# Gene-environment interactions using a Bayesian whole genome regression model

**DOI:** 10.1101/797829

**Authors:** Matthew Kerin, Jonathan Marchini

## Abstract

The contribution of gene-environment (GxE) interactions for many human traits and diseases is poorly characterised. We propose a Bayesian whole genome regression model, LEMMA, for joint modeling of main genetic effects and gene-environment interactions in large scale datasets such as the UK Biobank, where many environmental variables have been measured. The method estimates a linear combination of environmental variables, called an environmental score (ES), that interacts with genetic markers throughout the genome, and provides a readily interpretable way to examine the combined effect of many environmental variables. The ES can be used both to estimate the proportion of phenotypic variance attributable to GxE effects, and also to test for GxE effects at genetic variants across the genome. GxE effects can induce heteroscedasticity in quantitative traits and LEMMA accounts for this using robust standard error estimates when testing for GxE effects. When applied to body mass index, systolic, diastolic and pulse pressure in the UK Biobank we estimate that 9.3%, 3.9%, 1.6% and 12.5% of phenotypic variance is explained by GxE interactions, and that low frequency variants explain most of this variance. We also identify 3 loci that interact with the estimated environmental scores (− log_10_ *p* > 7.3).

## Introduction

Despite long standing interest in gene-by-environment (GxE) interactions^1^, this facet of genetic architecture remains poorly characterized in humans. Detection of GxE interactions is inherently more difficult than finding additive genetics in genome wide association studies (GWAS). One difficulty is that of sample size; a commonly cited rule of thumb suggests that detection of interaction effects requires a sample size at least four times larger than that required to detect a main effect of comparable effect size ^2^. Another is that an individual’s environment, which evolves through time, is very hard to measure in a comprehensive way, and is inherently high dimensional. Also, there are many environmental variables that could plausibly interact with the genome and many ways to combine them, and typically these factors are not all present in the same dataset. The recently released UK Biobank dataset, a large population cohort study with deep genotyping and sequencing, and extensive phenotyping^3^, offers a unique opportunity to explore GxE effects ^4–10^.

Models that consider environmental variables jointly can be advantageous, particularly if several environmental variables drive interactions at individual loci, or if an unobserved environment driving interactions is better reflected by a combination of observed environments. StructLMM^7^ models the environmental similarity between individuals (over multiple environments) as a random effect, and then tests each SNP independently for GxE interactions. However, StructLMM is not a whole genome regression model, so does not account for the genome wide contribution of all other variants, which is often a major component of phenotypic variance.

Advances in methods applied to detect genetic main effects in standard GWAS have shown that linear mixed models (LMMs) can reduce false positive associations due to population structure, and improve power by implicitly conditioning on other loci across the genome ^11–13^. Often these methods model the unobserved polygenic contribution as a multivariate Gaussian with co-variance structure proportional to a genetic relationship matrix (GRM) ^14–16^. This approach is mathematically equivalent to a whole genome regression (WGR) model with a Gaussian prior over SNP effect sizes ^11^. More flexible approaches have been proposed in both the animal breeding ^17, 18^ and human literature ^19–21^ to allow different prior distributions that better capture SNPs of small and large effects. The BOLT-LMM method^13^ uses a mixture of Gaussians (MoG) prior and shows this can increase power to detect associated loci in some (but not all) complex traits.

Here we present a new method called Linear Environment Mixed Model Analysis (LEMMA) which aims to combine the advantages of WGR and modeling GxE with multiple environments, and is applicable to large datasets with hundreds of thousands of individuals, such as the UK Biobank. Instead of assuming that the GxE effect over multiple environments is independent at each variant, as StructLMM does, we learn a single linear combination of environmental variables (that we call an environmental score (ES)), that has a common role in interaction effects genome wide. The ES is estimated within a Bayesian WGR model that uses two separate MoG priors on main genetic effects and GxE effects. We use variational inference to fit the model that is tractable for GxE analyses of biobank scale datasets with tens of environmental variables (see **Online Methods**).

Estimating the ES satisfies one of the primary motivations of this work, by providing a readily interpretable way to examine the combined effect of many environmental variables and how they might interact with genotype. A motivating example is the investigation of how modern obesogenic environments might accentuate the genetic risk of obesity. Tyrell et al.^22^ studied environments one at a time for their interaction with a body mass index (BMI) genetic risk score (GRS) and found several significant interactions. Our new method allows joint analysis of environments that might plausibly better represent an obesogenic environment, negating the need to model each environment one at a time. Our other motivations when developing LEMMA were to develop a powerful method to detect GxE interactions, and to estimate the proportion of variance that could be attributable to GxE interactions.

A LEMMA analysis has several distinct steps. First, the model is fitted using a large set of SNPs genome-wide, for example all the SNPs that have been directly assayed on a genotyping chip. The estimated ES is then used to estimate the proportion of phenotypic variability that is explained by interactions with this ES (GxE heritability), using randomized Haseman-Elston (RHE) regression^23, 24^. This heritability analysis can be run on genotyped or imputed SNPs, and can be stratified by minor allele frequency (MAF) and linkage disequilirium (LD) to better interrogate the genetic architecture of GxE interactions. The ES is also used to test for GxE interactions one variant at a time, typically at a larger set of imputed SNPs in the dataset. We use “robust” standard errors when testing each variant for a GxE interaction, which helps to control for the conditional heteroskedasticity caused by GxE interactions. We also suggest checks and solutions for the situation where environmental variables are themselves heritable and have a non-linear relationship to the trait of interest (see **Online Methods**).

We compared LEMMA to existing approaches such as StructLMM and F-tests using simulated data, and applied the approach to UK Biobank data for body mass index (BMI), systolic blood pressure (SBP), diastolic blood pressure (DBP) and pulse pressure (PP).

## Results

### Performance on simulated data Figure 1

compares the ability of different methods to detect GxE interactions at SNPs in simulations where a single true ES interacts with SNPs across the genome. **Figure S1** shows the false positive rate (FPR) to detect main effects. We compared our default version of LEMMA, which uses robust standard errors, StructLMM, a simple F-test of interaction and an F-test that uses robust standard errors (see **Online Methods**). The simulations vary GxE heritability, the total number of environmental variables and sample size. When sample size is large (N=100k), all the methods have reasonable control of FPR and LEMMA controls FPR at least as well as other methods across the range of simulations. When sample size is smaller (N=25k) the robust F-test performs less well as the number of environments grows (**Figure 1a**) and the F-test and StructLMM perform less well as the amount of GxE variance increases (**Figure 1b**). When we increase the sample size to N=200k we still find that LEMMA has a slighty inflated Type I error rate (see **Figure S2**).

**Figure 1:**
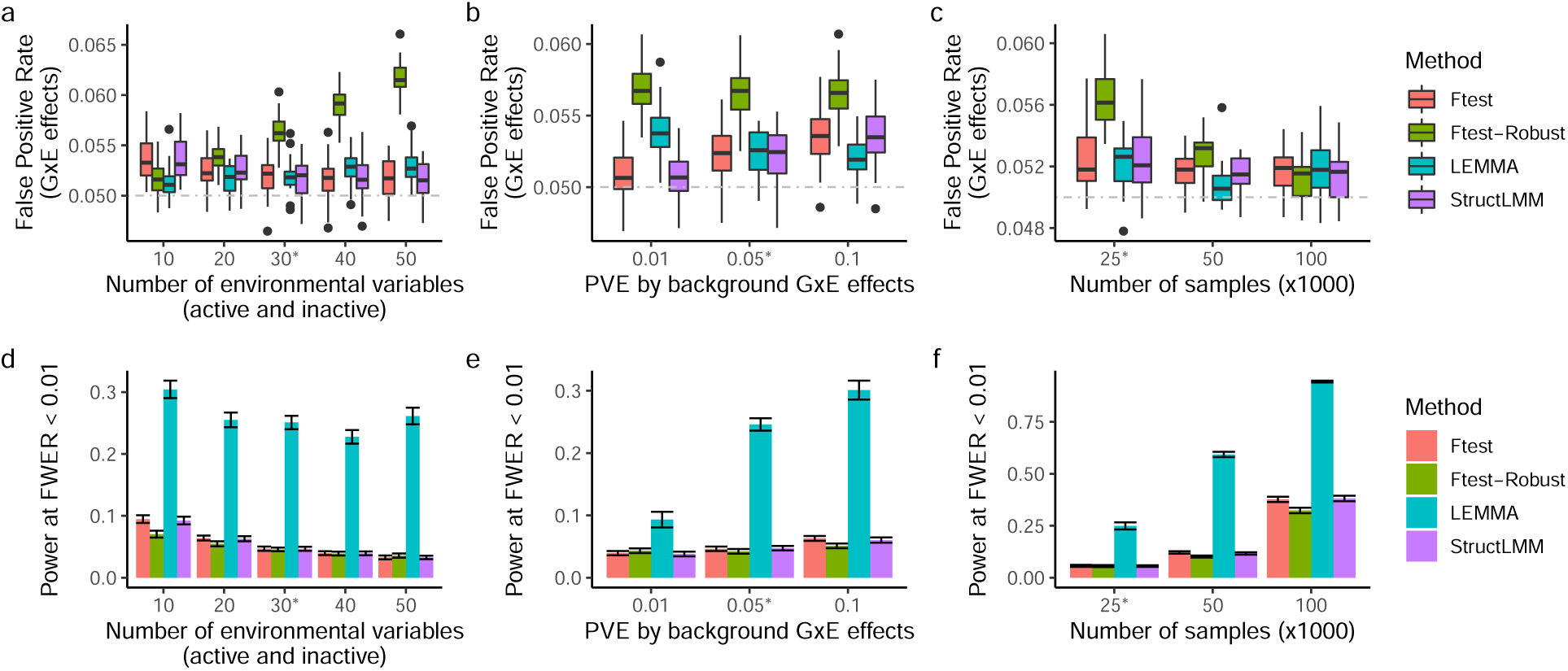
Type I error and Power of tests to detect GxE effects in simulation. (a-c) Comparison of false positive rate as the number of environments increases (a), as phenotype variance explained by GxE effects increases (b) or as the number samples increases (c). (d-f) Analogous comparison of the power to detect GxE interactions. Simulations used genotypes subsampled from the UK Biobank and by default contained *N* = 25*K* samples, *M* = 100*K* SNPs, 6 environmental variables that contributed to the ES and 24 that did not (default parameters denoted by stars). We assess power (at Family Wise Error Rate; FWER < 0.01) to detect 60 causal SNPs whose GxE effect each explained 0.00016% of trait variance. See **Online methods** for full details of phenotype construction.

It is interesting that all the methods we tested have slightly inflated Type I error, and this is likely due to a number of different reasons. StructLMM and the F-test fit a model at each variant and ignore GxE effects at other loci, which can induce heteroscedasticity that can inflate Type I error^49, 61^. We used robust standard errors for the robust F-test but it seems that this approximation works best when the number of environmental traits is small. LEMMA does account for GxE effects at other loci and also uses robust standard errors, but still has slightly inflated Type I error that gets worse as the number of environments increases (**Figure 1a** and **Figure S2**). In parallel simulations (see **Figure S3**) we find that our model slightly over-estimates GxE heritability as number of environments increases. Since our simulations test for GxE effects at SNPs used to estimate the ES we suspect that the Type I error inflation is due to this two stage approach.

When there is a single true ES involved in GxE interactions we found that LEMMA provided a substantial power increase (**Figure 1, Figure S4**). StructLMM and F-tests have very similar power in these simulations, although previous work suggests that StructLMM may outperform the F-test in small samples ^7^.

When estimating the GxE heritability of the LEMMA ES using randomized HE regression (RHE) with a single SNP component (RHE-SC) we observed some upward bias as the number of environments increases. This effect is ameliorated by increasing sample size (see **Figure S3**), suggesting that the influence of over-fitting in our Biobank analyses is mild. In twenty simulations with *L* = 30 environmental variables, *N* = 100k samples and true GxE heritability of 5% we observed mean GxE heritability of 5.2%. **Figure S5** further illustrates the ES estimation accuracy of LEMMA.

Finally, we ran LEMMA on two sets of simulated datasets (N=25k) with causal SNPs chosen either randomly, or chosen to be low frequency (MAF<0.1). We used the ES estimated from each simulated dataset to estimate 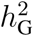 and 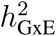 using RHE, with SNPs stratified by MAF and LD (RHE-LDMS), and then without any stratification (RHE-SC). Previous studies have established that estimating heritability using a single SNP component makes assumptions about the relationship between MAF, LD and trait architecture that may not hold up in practice^27, 28^, whereas stratifying SNPs into bins according to MAF and LDscore (the LDMS approach) is relatively unbiased^28–30^. **Figure S6** confirms that stratifying by MAF and LD results in accurate heritability estimates irrespective of the MAF distribution of causal SNPs, and suggests that this method can be used to interrogate the MAF distribution of GxE component of a trait using LEMMA. However when causal SNPs are low frequency, not stratifying by MAF and LD results in underestimation of 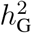.

### Controlling for heritable environmental variables

Previous work by Tchetgen *et al*.^25^ has shown that misspecifying the functional form of an environmental variable can induce heteroskedasticity into tests for GxE interactions. The authors further show that use of robust standard errors will control for this heteroskedasticity, but only if the environment is independent of the variant being tested. Independence between genotypes and the misspecified environment is important because it means that the (least squares) mean estimator is still unbiased.

However, environmental variables themselves often have a genetic basis. We therefore performed simulations where the phenotype depended on the non-linear (squared) effect of a heritable environmental variable. In simulation (**Figure 2a,c**) we observed that misspecification of the environmental variable can cause substantial inflation in GxE test statistics at heritable sites of the confounding environment. Relatively smooth non-linearities, such as squared effects, are easily detected by regression modeling before using LEMMA (see **Online Methods**) and can then be included as covariates (indicated in **Figure 2** by +SQE). This procedure produced well calibrated test statistics for all methods in simulation (**Figure 2c**).

**Figure 2:**
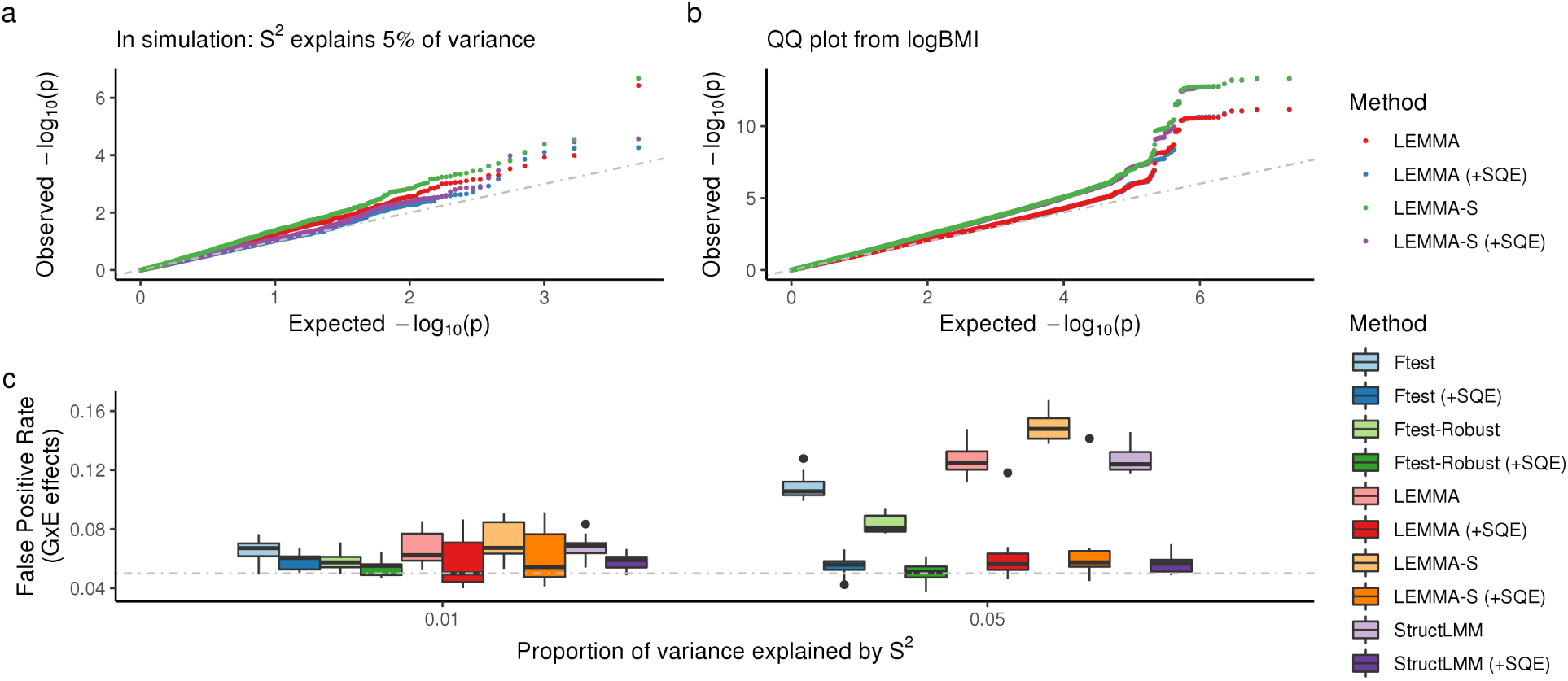
Bias from model mis-specification of a heritable environmental variable. (a) Comparison of GxE association test statistics from a single simulation where non-linear dependence on the confounder explains 5% of trait variance. FPR at heritable sites of the misspecified environment only. (b) Comparison of GxE association test statistics from an analysis of logBMI in 281, 149 participants from the UK Biobank. (c) False Positive Rate at heritable sites of the misspecified environment whilst the strength of squared dependence varies. 20 repeats per scenario. Abbreviations are as follows: LEMMA-S, LEMMA with non-robust variances used to compute test statistics; (+SQE), significant squared environmental variables (Bonferroni correction) included as additional covariates.

In **Figure 2b** we compare the GxE association test statistics from our analysis of logBMI in the UK Biobank, with and without adjusting for detected squared effects. Although we detected squared effects for 30 of the 42 environmental variables (significance level 0.01; Bonferroni correction for multiple testing), the ES obtained from the two analyses was almost identical (Pearson *r*^2^ > 0.999). As the additional variance explained collectively by the squared effects was negligible (incremental *R*^2^ < 0.00001) it would be surprising if this was not the case. Negative log_10_(*P*)-values from the two analyses were also highly correlated (Pearson *r*^2^ = 0.961), although there were small changes in the *p*-values at the *FOXO3* locus (which remained genome-wide significant in both analyses) and at the *SNAP25* locus (which was genome wide significant in the (-SQE) analysis only). We therefore conclude that the influence from this form of confounding in our analysis of logBMI was minor. However as the cost to this procedure is small, LEMMA uses the (+SQE) strategy by default for all analyses of UK Biobank traits.

### GxE interaction analysis in the UK Biobank

We applied LEMMA to characterize GxE interactions in Body Mass Index (logBMI), Systolic Blood Pressure (SBP), Diastolic Blood Pressue (DBP) and Pulse Pressure (PP) using a set of 42 environmental variables similar to those used in previous analyses ^7, 8, 26^, including data on smoking, hours of TV watched, Townsend Index, physical exercise and alcohol consumption (see **Online methods** and **Table S1)**.

We analyzed GxE heritability due to multiplicative effects with the ES using both *M* = 639, 005 genotyped SNPs and *M* = 10, 270, 052 common imputed SNPs (MAF ≥ 0.01 in the full UK Biobank cohort), stratified by MAF and LDscore into 20 components. Using imputed SNPs we estimated GxE heritability of 9.3%, 12.5%, 3.9% and 1.6% for logBMI, PP, SBP and DBP respectively (see **Table 1**). On genotyped SNPs the GxE heritability estimates were slightly lower for logBMI and PP (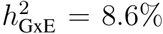 and 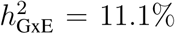 respectively) and almost identical for SBP and DBP (see **Table S2**). For all traits the heritability of additive SNP effects were slightly higher on imputed data, consistent with previous results ^29^.

**Table 1:** Partitioned heritability estimates for four quantitative traits in the UK Biobank. Heritability estimates obtained using common imputed SNPs (MAF> 0.01 in the full UK Biobank cohort) with RHE-LDMS. GxE heritability estimates were were obtained using the ES from each model fit. All analyses controlled for the same covariates used in the WGR analysis (including the top 20 principal components). Abbreviations; s.e, standard error estimated using the block jackknife (see **Online Methods**); 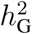, heritability due to additive genetic effects; 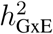, heritability due to multiplicative GxE effects; RHE, randomized HE-regression^23, 24^; LDMS, SNPs stratified by minor allele frequency and LDscore (20 components); INT, Inverse normal transform applied to males and females separately.

| Trait | $h_G^2$ (s.e) | $h_{G \times E}^2$ (s.e) |
| --- | --- | --- |
| logBMI | 0.274 (0.056) | 0.093 (0.028) |
| INT(BMI) | 0.278 (0.056) | 0.059 (0.024) |
| BMI | 0.268 (0.055) | 0.137 (0.031) |
| PP | 0.228 (0.051) | 0.125 (0.028) |
| SBP | 0.251 (0.05) | 0.039 (0.023) |
| DBP | 0.254 (0.05) | 0.016 (0.02) |

When working with quantitative traits it can be hard to choose an optimal transformation or scale for each trait. Tyrell et al. ^22^ analyzed BMI using the raw scale and then also by transforming to a standard normal distribution. They observed larger interaction effects on the raw scale and suggested that this was due to larger variance in BMI in individuals in the high-risk environment groups, which causes heteroscedasticity, and inflates effect estimates. In addition to our main analysis, which used log BMI, we re-ran LEMMA using the raw BMI measurement, and then also by transforming to a standard normal distribution in females and males separately. These results are presented in **Table 1** and agree with the results of Tyrell et al. ^22^, with estimates of GxE heritability on the raw, log and inverse normal scale of 13.7%, 9.3% and 5.9% respectively.

Previous work on models of natural selection has suggested that the variance explained by additive SNP effects should be uniformly distributed as a function of MAF in a neutral evolutionary setting^31^, and that enrichment of the variance explained by low frequency SNPs is evidence for negative selection. For all four traits we found that variance explained by the additive genetic effects of low frequency SNPs (MAF < 0.1) was slightly elevated, consistent with previous observations of negative selection^32^ (**Figure 3**). Additionally the distribution of additive genetic effects by MAF for logBMI was qualitatively similar to that found by GREML-LDMS is a previous study^29^. In contrast we found that variance explained by GxE effects was overwhelming attributed to low frequency SNPs (MAF < 0.01), especially those with low LD. However we are not aware of any evolutionary theory that has been extended to model the MAF distribution of GxE effects.

**Figure 3:**
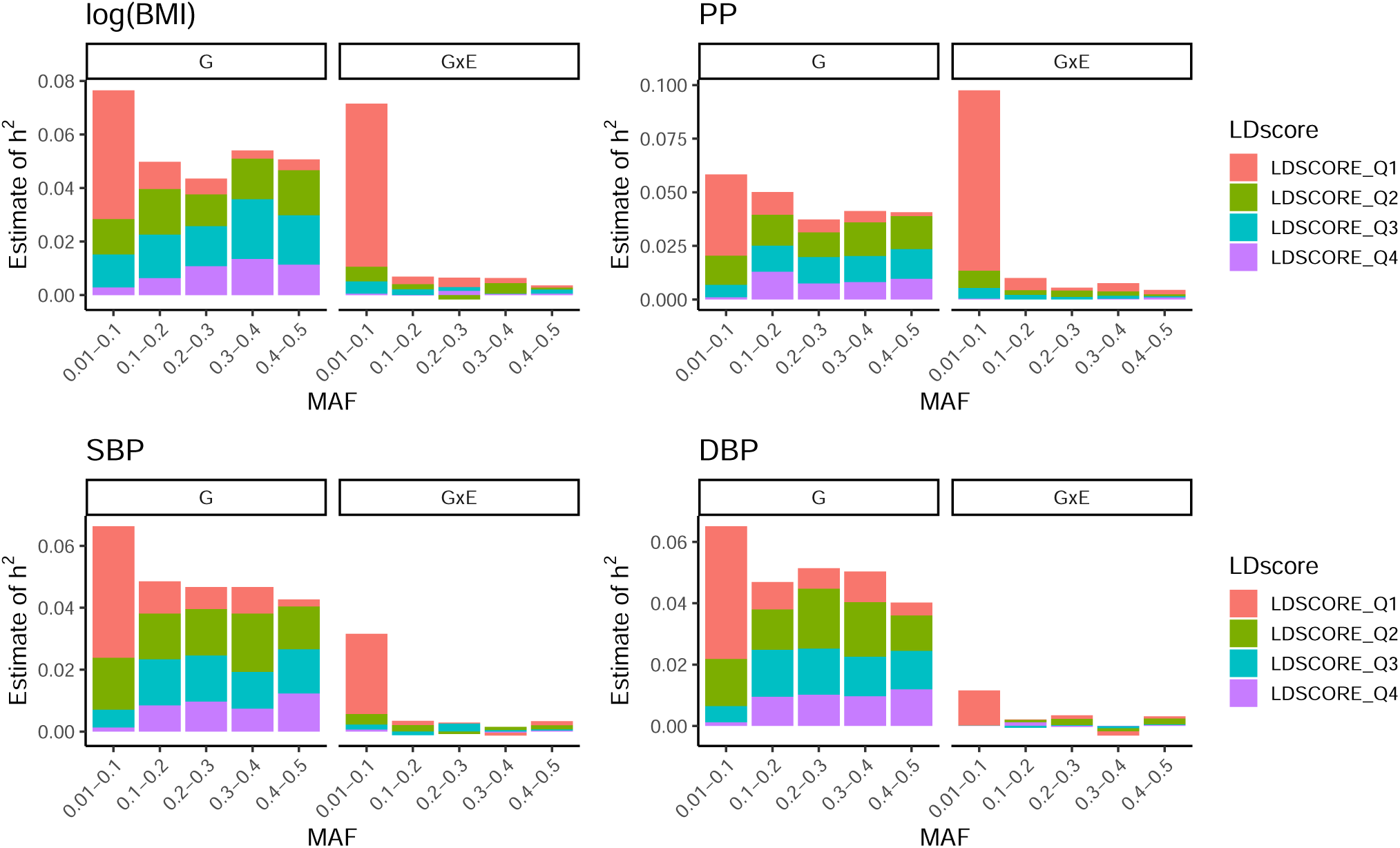
Partitioned heritability estimates for four quantitative traits in the UK Biobank. Heritability estimates partitioned into additive genetic and multiplicative GxE interaction effects for four quantitative traits in the UK Biobank, using approximately 280, 000 unrelated white British individuals (see **Table S1**) and *M* = 10, 270, 052 common imputed SNPs (MAF > 0.01 in the full UK Biobank cohort). Multiplicative GxE interactions were computed using the *ES* from each model fit. Heritability estimation was performed using a multi-component implementation of randomized HE-regression^23, 24^ with SNPs stratified into 20 components (5 MAF bins and 4 LD score quantiles).

For logBMI we estimated an ES that put high weight on alcohol intake frequency, Townsend Index and physical activity measures (**Figure 4c**). Almost all of the non-dietary environmental exposures had a higher effect in women than in men, with smoking status being the one exception. This is reflected in the facts that (a) the ES having much higher variance in women and (b) those with a negative ES were almost all female (97%) (see **Figure 4b**). When comparing the characteristics of those in the bottom 5% of the ES to the whole cohort (using the mean for continuous variables and the mode for categorical), we found that those in the bottom 5% were predominantly female (100% vs 53%), younger (51 vs 56), had higher Townsend deprivation index (0.91 vs −1.74), drank less often (‘Special occasions’ vs ‘Once or twice a week’) and watched more TV (3.28 vs 2.69 daily hours of TV) (**Table S3**). We note that positive values of the Townsend index indicate material deprivation, whereas negative values indicate relative affluence.

**Figure 4:**
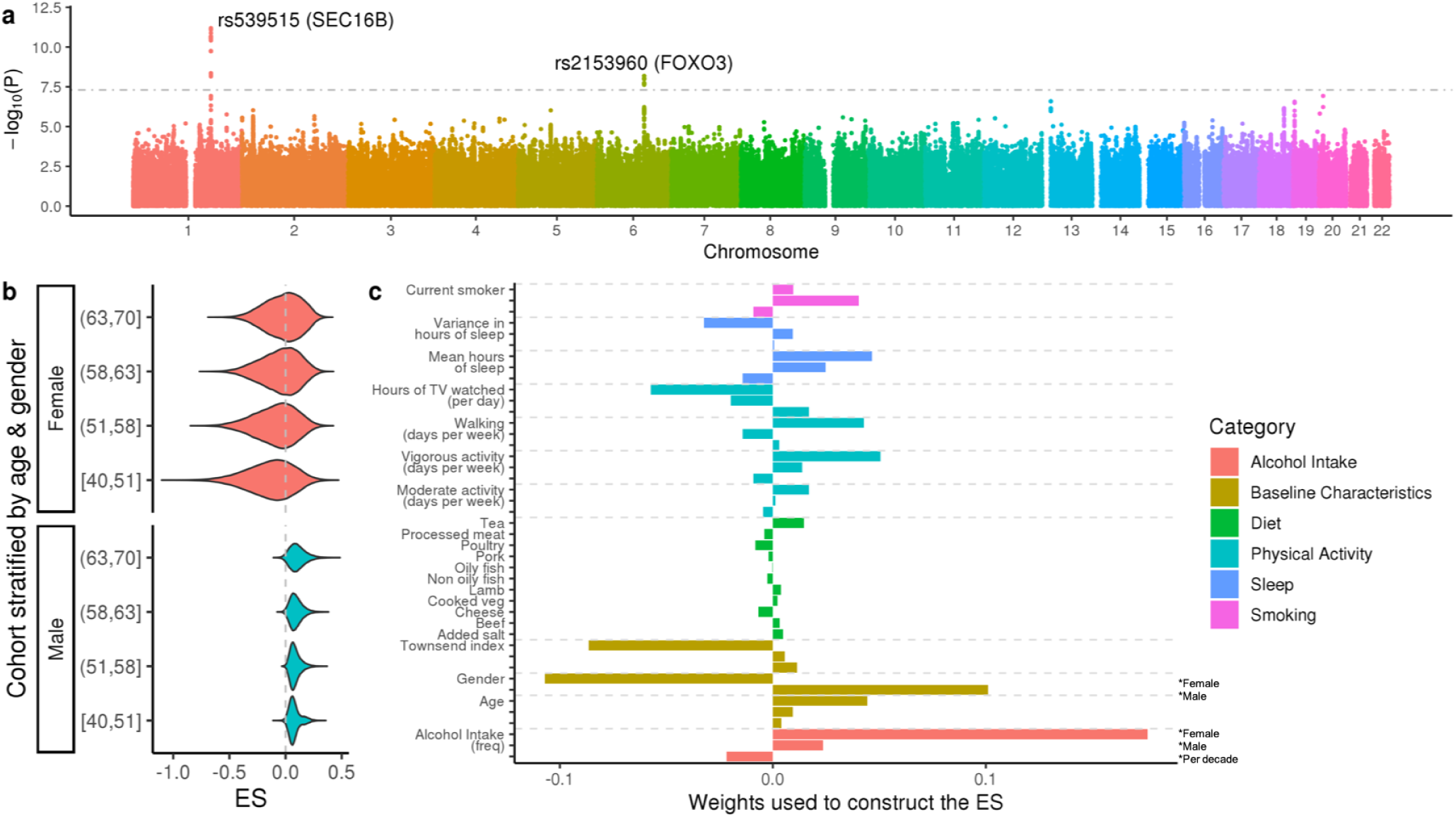
GxE analysis of logBMI in the UK Biobank. (a) LEMMA association statistics testing for multiplicative GxE interactions at each SNP. The horizontal grey line denotes (*p* = 5 × 10^−8^), *p*-values are shown on the − log_10_ scale. (b) Distribution of the environmental score (ES), stratified by gender and age quantile. (c) Weights used to construct the ES. Dietary variables have a single weight shown on the per standard deviation (s.d) scale. ‘Gender’ has two weights; a gender specific intercept for women (first) and men (second). Remaining non-dietary variables have three weights; (first) a per s.d effect for women only, (second) a per s.d effect for men only, (third) a per s.d per decade effect which is the same for both genders. s.d for the male and female specific weights is computed for each gender separately. Age is computed as the number of decades aged from 40. See **Online Methods** for details.

Previous cross-sectional studies have reported GxE interactions between a linear predictor formed from BMI-associated SNPs and alcohol intake frequency ^33^, Townsend Index ^22, 33^, physical activity measures ^5, 22, 33, 34^ and time watching TV ^22, 33, 34^, all of which had high relative weight in the logBMI ES. An alternative approach from Robinson *et al*.^6^ binned samples according to their environmental exposure (eg. age) and tested for significant differences in SNP heritability using a likelihood ratio test. They reported strong interaction effects with age in a cohort of 43, 407 individuals whose ages spanned 18 − 80, but only reported significant interactions with smoking in the UK Biobank interim release. This suggests that we might expect age to play a more dominant role in the logBMI ES in a cohort that included younger individuals. Finally, one category that is notably down weighted is the contribution from dietary variables. Although significant interactions with fried food consumption ^35^ and sugar sweetened drinks ^36^ have previously been reported in a cohort of US health professionals, these dietary variables were not included in the diet questionnaire used by the UK Biobank.

The ES for PP was dominated by the effects of age and gender (age, age^2^, age-x-gender, gender together explained 94.9% of variance in the ES). The magnitude of the ES was strongly associated with increased age^37^ whilst the sign of the ES was strongly associated with gender, implying that GxE effects were stronger in the elderly but acted in the opposite direction in men and women (**Figure S7**).

Similarly we observed that variance of the ES increased with age in both SBP and DBP, but instead of age itself being highly weighted we found that age interactions with other environmental variables were most important for explaining variation in the ES. Specifically for SBP we found that age interactions with smoking, Townsend index and alcohol frequency explained 86% of variance in the ES (**Figure S8**) When compared to the cohort average, we found that participants in the top 5% of the SBP ES were older (63 vs 58), had higher Townsend deprivation index (1.2 vs −1.74) and were more likely to smoke (59% vs 9%) whereas those in the bottom 5% were also older (65 vs 58), predominantly female (91.5%), rarely drank alcohol (43.9% drank “Never”) and had low Townsend deprivation index (−2.9 vs −1.74) (**Table S4**).

Finally we observed notably higher variance in the ES for DBP among men, most of which appeared to be driven by high gender-specific weights for smoking status and alcohol frequency (**Figure S9**). We further observed that alcohol frequency and smoking status became increasingly influential with age. The total SNP-GxE heritability for this ES however was quite low.

When testing for significant GxE interactions between the estimated ESs and imputed markers across the genome we observed that use of the robust standard errors made a noticeable difference to the calibration of LEMMA (**Figure S10**; **Table S5**). We identified two loci for logBMI (**Figure 4a**), one locus for DBP (**Figure S9a**) and zero loci for SBP and PP, using a threshold of 5 × 10^−8^ for genome wide significance (**Table 2**). This table also includes results from a standard linear regression GWAS test at the 3 loci (see also **Table S6**).

**Table 2:**
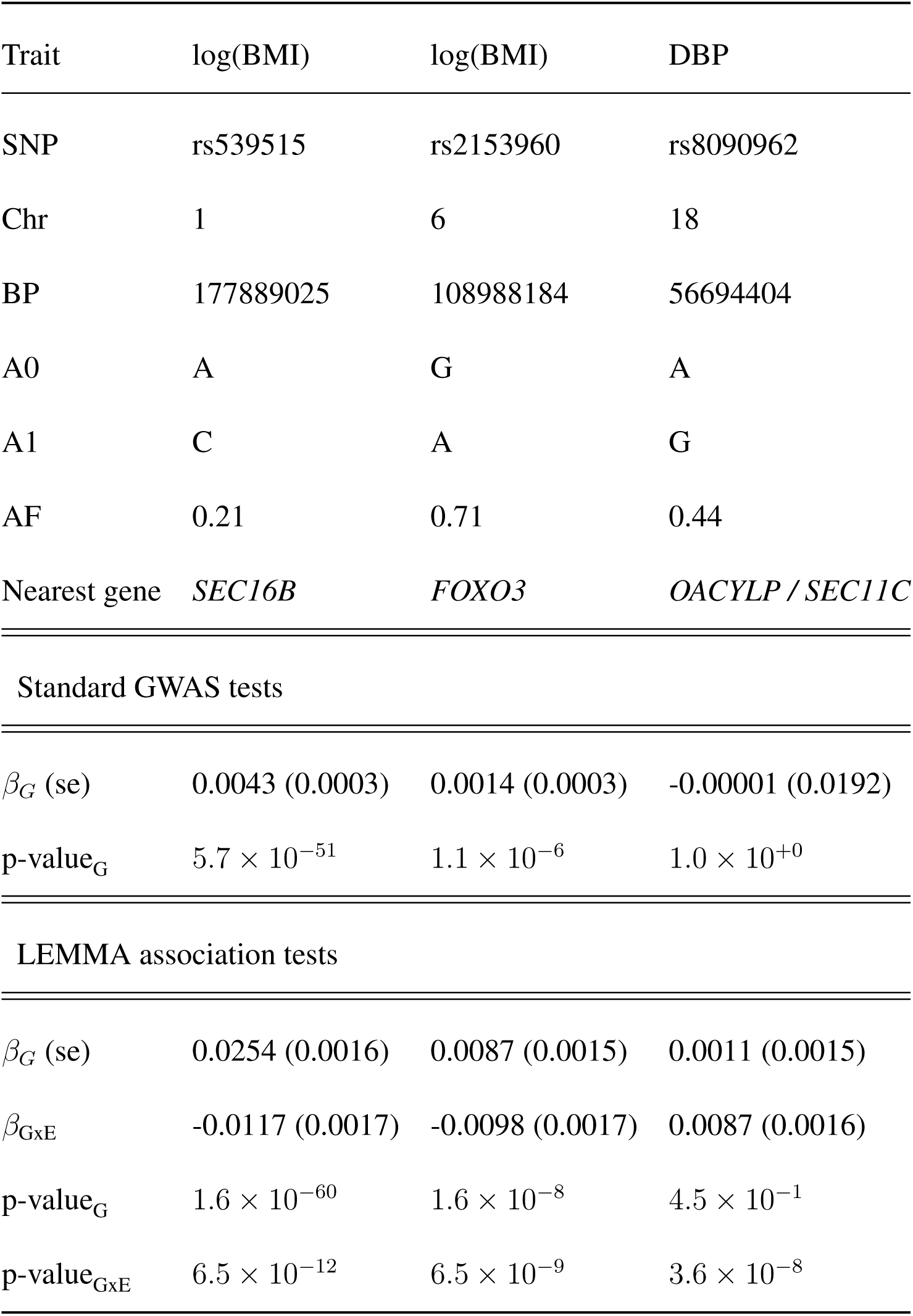
Loci with genome-wide significant GxE interaction effects with the ES. Independent loci with genome-wide significant (*P* < 5 *×* 10^−8^) GxE interaction effects with the environmental score (ES). Loci at least 0.5cM apart were judged to be independent. SNP effect sizes reported on a per s.d scale. SNP locations follow the GrCh37 human genome assembly. All loci had IMPUTE INFO score > 0.99. Abbreviations are as follows; A0, reference allele; A1, alternative allele; AF, reference allele frequency; s.d, standard deviation.

For logBMI, LEMMA identified GxE interactions at rs2153960 (*p* = 6.5×10^−9^; **Figure S11**) and at rs539515 (*p* = 6.5 × 10^−12^; **Figure S12**). The SNP rs2153960 is an intron in the *FOXO3* gene and has been previously associated with Insulin-like growth factor 1 (IGF-I) concentration in a cohort of 10, 000 middle aged Europeans ^38^. IGF-I is known to be a a central mediator of metabolic, endocrine and anabolic effects of growth hormone and is also involved in carbohydrate homeostasis ^38^. The patterns of main effect association and GxE association show considerable overlap (**Figure S11a**). This SNP did not reach genome-wide levels of significance using the standard linear regression GWAS test (**Table 2**).

The SNP rs539515 is located 6kb downstream of *SEC16B*. The patterns of main effect association and GxE association are very similar **Figure S12a**. Multiplicative GxE interactions have been reported at *SEC16B* with multiple environmental variables in a similar analysis in the UK Biobank^7^, and with physical activity separately in Europeans ^5^ (*p* = 0.025) and in Hispanics ^39^ (*p* = 8.1 × 10^−5^). Highly significant variance effects (*p* = 3.88 × 10^−17^), which can be indicative of GxE, have also been reported at the *SEC16B* locus using *N* = 456, 422 Europeans in the UK Biobank^40^. GxE interactions have been reported at *SEC16B* with multiple environmental variables in a similar analysis in the UK Biobank^7^ and with physical activity separately in Europeans ^5^ (*p* = 0.025) and in Hispanics ^39^ (*p* = 8.1 × 10^−5^). *SEC16B* transcribes one of the two mammalian homologues of the SEC16 protein, which has a key role in organizing endoplasmic reticulum exit sites by interacting with COPII components ^41^. Although several GWASs have identified associations between *SEC16B* ^42, 43^, the relevance of *SEC16B* to BMI is not well characterized ^44^. Some evidence exists to suggest that *SEC16B* has role in the transport of peroxisome biogenesis factors; peroxisomes being an organelle involved in the catabolism of long chain fatty acids found ubiquitously in eukaryotic cells. Previous authors^43^ have also speculated that the *SEC16B* might play a role in the transport of appetite regulatory peptides, however we are not aware of any evidence for this theory.

The DBP associated SNP is rs8090962, but only just passes our threshold for significance and we are least confident that this is a true GxE association for a few reasons. The SNP is located within an enhancer, approximately 100KB downstream of the *SEC11C* gene and 50KB upstream of *ZNF532*. Neither gene has previously been associated with blood pressure traits. There is some evidence of a main effect close by (**Figure S13**a) but the pattern of main effect association does not coincide well with the pattern of GxE associations. In addition the pattern of GxE association by genotype (**Figure S13**b) shows a striking cross over by genotype between extremes of the ES. We have observed above that our test statistics is very slightly inflated so this could be a false positive association.

### Relationship of genetics PCs and environmental scores

We regressed the estimated ESs against the PCs for each of the 4 UKB traits and the results are included in **Table S7**. We found some significant associations, mostly with PC5, which seems to correlate with North-South geography in the UK^45^. To explore further we also re-ran the heritability analysis by including interaction terms of the ES with the genetic PCs as control variables, but the results were almost unchanged (see **Table S8**).

### Comparison of the LEMMA ES with a marginal ES

For each trait, we used least squares regression to compute a linear model fit using all of the non genetic covariates used in the LEMMA analysis. We then constructed an environmental score (referred to as ES_marginal_) using the marginal environmental effects from this model fit. The correlation between the LEMMA ES and ES_marginal_ was −0.062, −0.019, −0.297 and −0.088 for logBMI, PP, SBP and DBP respectively, suggesting that these vectors are quite dissimilar. **Figure S14** shows a comparison of the interaction weights used to construct the LEMMA ES and ES_marginal_ for each of the four traits. Visually the weights learnt through each approach look quite distinct. In particular, age, age^2^ and age×gender have much higher relative weight in ES_marginal_ than in the LEMMA ES.

### Comparison of methods on UK Biobank data

To compare LEMMA with existing single SNP methods we also ran StructLMM, the F-test and the robust F-test on logBMI using the same set of environmental variables as used by LEMMA (but not including the significant squared environments as covariates). Manhattan plots are displayed in **Figure S15**. Test statistics from both the F-test (*λ*_GC_ = 1.37) and StructLMM (*λ*_GC_ = 1.235) were substantially inflated when compared to the robust F-test and LEMMA (*λ*_GC_ = 1.03 and *λ*_GC_ = 1.062 respectively; see **Table S5**), suggesting that StructLMM does not properly control for heteroskedasticity. There are clear differences between the 4 methods, especially among SNPs with suggestive evidence of GxE Interaction results (**Figure S16a**). LEMMA did not find the FTO locus, StructLMM and F-test did not find the SEC16B locus, and the robust F-test only found the FTO locus.

LEMMA relies on the assumption that all GxE interaction effects for a single trait share a common ES, and we have shown in simulation that when this assumption holds LEMMA achieves substantial increases in power. However we would expect LEMMA to have little power to detect SNPS which interact with a combination of environments that is not well correlated with the genome-wide ES estimated by LEMMA. The *FTO* seems to be one clear example of this. We extracted an estimate of the SNP specific interaction profile at *FTO* using the robust F-test (**Online Methods**), we found that it’s correlation with LEMMA’s ES was low (Pearson *r*^2^ = 0.3). In comparison, a similar analysis at *SEC16B* and *FOXO3* yielded much higher correlations (Pearson *r*^2^ = 0.725 and *r*^2^ = 0.713 respectively).

## Discussion

In this study we proposed a new whole genome regression method, LEMMA, that estimates a single environmental score (ES) that interacts with SNPs across the genome. In simulation we have demonstrated that the ES can be used to compute well calibrated p-values of the multiplicative interaction effect at each SNP. LEMMA is also able to quantify the trait variance attributable to MAF and LD stratified interaction effects of the ES.

In analyses of four quantitative traits in the UK Biobank, we have demonstrated that GxE effects among common imputed SNPs make a non-trivial contribution to the heritability of logBMI and PP (9.3% and 12.5% respectively). Our stratified heritability analysis has suggested that GxE interactions for these traits are mostly driven by low frequency variants. Our analysis identified three loci with statistically significant GxE interaction effects. Two of these loci, rs539515 (*FOXO3*) and rs8090962, are novel and for the other, rs539515 (*SEC16B*), we show stronger evidence for than in the previous study^7^.

Robinson *et al*.^6^ have previously attempted to quantify the contribution of GxE interactions to the heritability of BMI in a study performed on imputed SNPs from the interim UK Biobank release. Using the GCI-GREML model implemented in GCTA^46^ and eight environmental variables that included measures of smoking, hours of TV watched and alcohol frequency, Robinson *et al*.^6^ reported that only smoking had significant GxE heritability (4.0%). In contrast, the ES estimated for logBMI in our analysis had non-zero contributions from many environmental variables, including hours of TV watched and smoking, suggesting that multiple environmental variables can influence on the genetic predisposition to BMI. Modeling these environmental variables jointly allowed LEMMA to capture a combination whose GxE interactions explained 9.3% of heritability.

We have also evaluated the performance of three existing single SNP methods (StructLMM, the F-test and a robust F-test), both in simulation and on logBMI from this same dataset. In simulation with large datasets we observed that StructLMM and the F-test had similar performance; an observation that also held in our analysis of logBMI. Both of these methods appeared vulnerable to heteroskedasticity, which we showed is likely to occur in traits with non-trivial GxE heritability. A simple adjustment, using ‘robust’ or Huber-White variance estimators, solved this problem. The two F-test methods further benefit from a wealth of existing theory^47^ and, being theoretically simpler than StructLMM, could be easily implemented as an R-plugin with PLINK^48^ (for example ^49^). In our opinion, the robust F-test is therefore the most appropriate of the three single SNP methods to model GxE effects with tens of environments in biobank scale datasets.

Although LEMMA represents a method with increased power to detect GxE interaction effects, our approach does have some caveats. First the gain in power is dependent on a strong assumption on the underlying genetic architecture. Whilst our analysis suggests that this does hold to some extent for PP and logBMI, this may not be the case for other traits.

In addition, LEMMA only estimates the proportion of phenotypic variance that is explained by interactions with this ES, and we do not claim that this captures all the GxE heritability of a trait. If relevant GxE environments are not included in the analysis, and these environments have low correlation to the environments that are included, then LEMMA cannot account for them, and will likely underestimate the true GxE heritability. Unobserved environments can cause trait variance to depend on genotype^8^ (see **Figure S19**) and extending LEMMA in this direction is left for future work.

LEMMA has the requirement that none of the environmental variables have any missing values. This could lead to a reduction in samples size if many environmental variables are included. If the amount of missing data is small it should not pose a big problem, and missing data imputation methods are also an option. If LEMMA is applied in situations where the missing data structure is related to the phenotype of interest then this could cause bias in the results.

Despite much effort to provide an efficient implementation, the LEMMA algorithm is still a computationally demanding. Using randomized HE-regression to estimate an improved initialization of the interactions weights may help to reduce run time, and is an avenue that we are currently pursuing.

Finally, for simplicity LEMMA currently searches only for GxE interactions with a single linear combination of environments. Generalizing the LEMMA approach to several orthogonal linear combinations or using functional annotation to restrict the SNPs that each ES interacts with, may yet yield more power to identify interactions in complex traits and explain more phenotypic variation.

## Online Methods

### Linear Environment Mixed Model Analysis (LEMMA)

The standard LMM used in genome wide association studies is written as

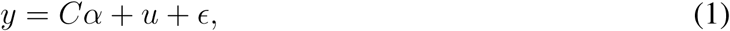

where *y* is the centered and scaled *N* × 1 vector of phenotypes, *C* is an *N* × *L*′ matrix of covariates with *L*′ × 1 fixed effects vector *α*, and *u* and *ϵ* are *N* × 1 vectors of unobserved polygenic and residual effects vectors respectively. Typically *u* is modeled as Gaussian with mean zero and covariance matrix 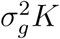. Specification of the *N* × *N* kinship matrix *K* is an area of active research ^50–53^ but the simplest approach is to let *K* = *XX*^*T*^ */M* where *X* is the *N* × *M* genotype matrix and columns of *X* (which usually correspond to SNPs) are normalized to have mean zero and variance one. This can equivalently be written as a Bayesian whole genome regression (WGR) model

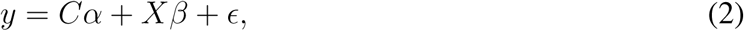

where

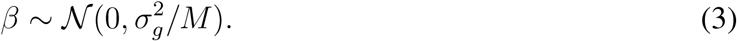

Here *β* is a *M* × 1 vector modeling the random effect of each SNP. This form corresponds to the so called infinitesimal model where every SNP is allowed to have a small but non-zero effect on a given trait. To generalize the model to a non-infinitesimal g enetic a rchitecture, w e model SNP effects with a mixture of Gaussians prior. This approach has been applied previously in human genetics ^13, 21^ and by the ‘Bayesian alphabet’ of genomic prediction methods in the animal breeding literature ^17, 18, 54^.

We extend this setup to model GxE interactions genome-wide with a linear combination of multiple environmental variables using

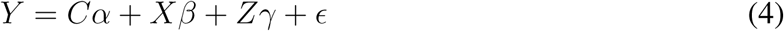

where

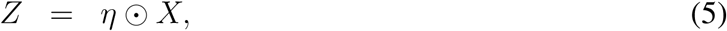

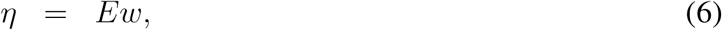

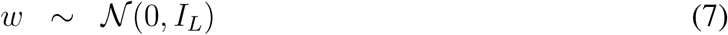

where *E* is an *N* × *L* matrix of environmental variables that could potentially be involved in GxE interactions and *w* is an *L* × 1 vector of weights. Nether they define the *N* × 1 vector *η* that is the linear combination of environments that we refer to as the environmental score (ES). This ES is learned in tandem with SNP effects. We note that all environmental variables contained in *E* must also be contained in *C*, so *L* ≤ *L*′. We chose to model the interaction weights *w* with a Gaussian prior, but in theory one could consider sparser priors such as a spike and slab. We set the variance of the prior on *w* to the identity matrix *I*_*L*_. Setting the prior variance of *w* to a parameter would be unidentifiable, as any change in scale would be absorbed by the prior variances on the interaction effects *γ* (see 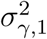 and 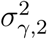 in equations 8-9).

The *N* × *M* matrix *Z* contains all of the multiplicative interaction terms of the ES *η* with all of the genetic variants. We use the notation *η* ⊙ *X* for the element-wise product of *η* with each column of *X*. In other words, *η* ⊙ *X* = diag (*η*) *X* where diag (*η*) is an *N* × *N* diagonal matrix with *η* as the diagonal. The vector of interaction effect sizes *γ* has dimension *M* × 1.

We chose to use MoG priors on both the main genetic effects (*β*) and the interaction effects (*γ*) as this prior is very flexible and spans the range of genetic architectures from polygenic to a very sparse model. The priors are

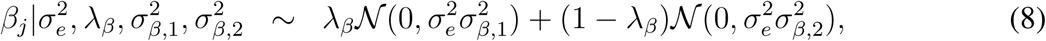

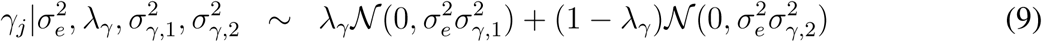

We use standard Gaussian priors on the covariate and error terms.

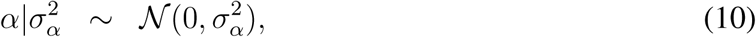

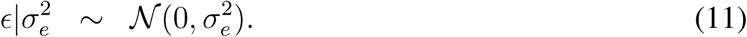

### Variational Inference

For notational convenience we define *θ* = {*α, β, γ, w*} as the set of latent variables, *𝒟* := {*X, E*} the genetic and environmental data and *ϕ* as the set of hyper parameters Then the posterior *p*(*θ*|*y, 𝒟, ϕ*) is given by

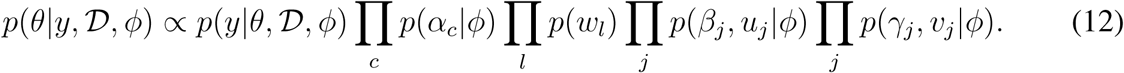

To evaluate the posterior we use the variational inference framework; approximating the true posterior *p*(*θ*|*y, 𝒟, ϕ*) with a tractable alternative distribution *q*(*θ*; *ν*) governed by (variational) parameters *ν*. To make inference tractable we use the standard Mean Field assumption so that *q*(*θ*; *ν*) factorizes

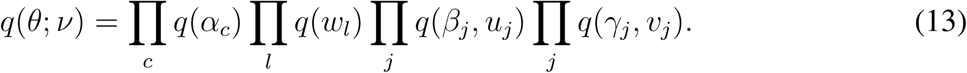

To make *q*(*θ*; *ν*) a close approximation of the true posterior we minimize the KL Divergence between *q*(*θ*; *ν*) and *p*(*θ*|*y, 𝒟, ϕ*) with respect to variational parameters *ν*. In this manner, the problem has been transformed from one of computing posterior distributions into one of optimization. We can show that minimizing the KL Divergence is equivalent to maximizing a lower bound on the marginal log likelihood by observing

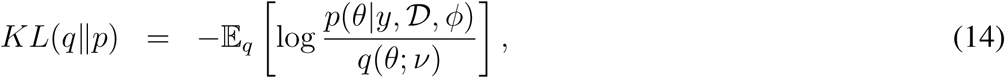

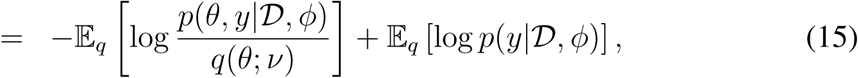

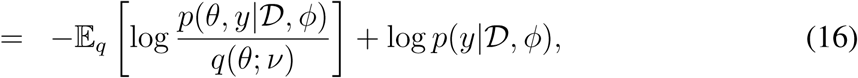

thus we can write

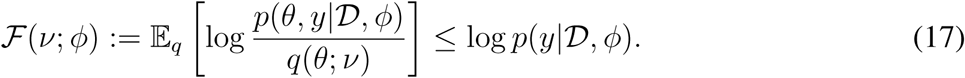

Here *ℱ s*(*ν*; *ϕ*) is commonly referred to as the Evidence Lower Bound (ELBO). Due to the factorized form of Equation (13) we can cyclically update the approximate distribution for each latent variable in turn until we reach convergence.

Our model depends on a set of eight hyper-parameters 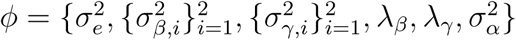. We set 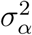 to a large constant to create a flat prior on the covariates, leaving seven unknowns. Similar methods have performed a grid search over hyper-parameter values (using either cross validation^13^ or the in-sample ELBO to identify the optimum^20^). For LEMMA a grid search would be computationally demanding, both because the set of hyper-parameters is larger and because we cannot efficiently perform multiple runs in parallel as done by Loh *et al*.^13^. Instead we maximize a lower bound on the approximate log likelihood (the so called Evidence Lower Bound or ELBO) with respect to the hyper-parameters. In this manner our approach can be viewed as a variational expectation maximization algorithm ^55, 56^.

Similar to the EM algorithm, the hyper-parameter maximization step can lead to slow exploration of the hyper-parameter space and thus to slow convergence of the LEMMA algorithm. We use an accelerator, SQUAREM ^57^, to speed up convergence. Given two estimates of the hyperparameters *ϕ*_*t*−2_ and *ϕ*_*t*−1_ we can adjust the maximized estimate *ϕ*_*t*_ with

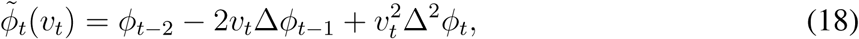

where Δ*ϕ*_*t*−1_ = *ϕ*_*t*−1_ − *ϕ*_*t*−2_ and Δ^2^*ϕ*_*t*_ = *ϕ*_*t*_ − 2*ϕ*_*t*−1_ + *ϕ*_*t*−2_. Thus the new adjusted estimate 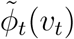 is a continuous function of the step size *v*_*t*_, which yields the original estimate *ϕ*_*t*_ for *v*_*t*_ = −1. As recommended by Varadhan *et al*.^57^ we set 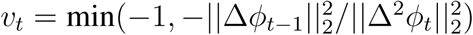. Occasionally this yields an estimate that is either outside of the domain of *ϕ* or leads to a state with worse ELBO than the previous state. For the first issue we use a simple backtracking method of halving the distance between *v*_*t*_ and −1, and for the second we simply judge model convergence when the absolute change in ELBO drops below a given threshold. We use the same convergence criterion as the BOLT-LMM method ^13^; namely that a full pass through all latent variables yields an absolute change of less than 0.01 in the approximate log-likelihood (ELBO). **Figure S17** shows the evolution of the ES parameter estimates for the four UK Biobank traits we analyzed, and illustrates that at the point of convergence the parameters appear stable.

### Identifying GxE associated loci

After convergence of the LEMMA variational inference algorithm, we obtain posterior mean estimates of 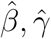 and 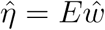. From these we construct residualized phenotypes following a Leave One Chromosome Out (LOCO) scheme;

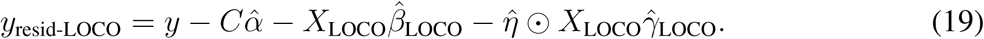

*X*_LOCO_ denotes the genotype matrix excluding SNPs on the same chromosome of the test SNP, and *β*_LOCO_ and *γ*_LOCO_ are constructed similarly. Using a LOCO scheme has been shown to increase power in LMMs as the effect of the test SNP is conditioned on the effects on a large proportion of the rest of the genome ^12, 15^.

For each imputed SNP, we then perform hypothesis tests *β*_test_ ≠ 0 and *γ*_test_ ≠ 0 using the linear model

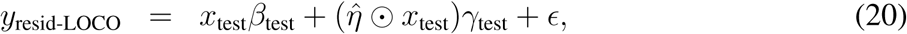

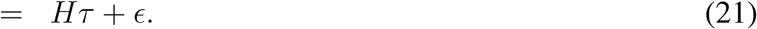

Here *H* is the *N* × 2 design matrix with columns containing *x*_test_ and 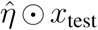 respectively, and *τ* is the 2 × 1 vector containing parameters *β*_test_ and *γ*_test_ respectively, which are the main genetic effect and interaction effect of the SNP being tested.

Assuming that *ϵ* has mean zero and covariance matrix Ω we can use the standard OLS estimator

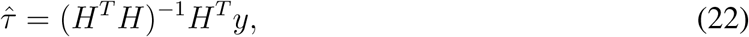

which (under certain regularity conditions) is asymptotically normally distributed with mean *τ* and variance Var 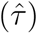. By assuming the residual phenotype is homoskedastic, that is that 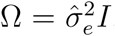, we can obtain the usual variance estimator given by

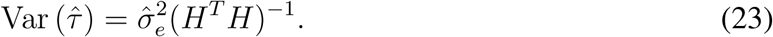

It has previously been observed that GxE interaction tests are likely to suffer from conditional heteroskedasticity^49^, and hence the homoskedastic variance estimator is likely to underestimate the true variance ^58^. We explain this phenomenon in detail in the **Supplementary Material**.

To overcome this we use robust standard errors, alternatively called Huber-White, sandwich or “heteroskedastic consistent” errors ^59, 60^, that are standard tools in economics ^47^ and have previously been proposed for use in GxE interaction studies ^25, 49, 61^. We further include a small adjustment that reduces bias in small samples ^62^. This yields the variance estimator

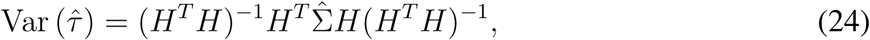

where 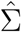 is a diagonal matrix with 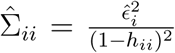, where 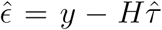 and *h* = *H*(*H*^*T*^ *H*)^−1^*H*^*T*^. Hence our GxE test statistic is given by

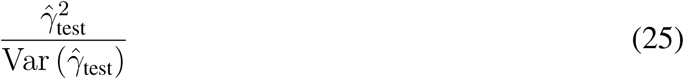

and under the null hypothesis is asymptotically distributed as 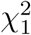. As main effects tests are not sensitive to assumptions of heteroskedasticity in the same way that GxE tests are ^49^, we use a simple t-test to test the hypothesis *β*_test_ ≠ 0.

### Heritability estimation

Previous genome wide regression methods ^20, 32, 63^ have shown that it is possible to rearrange the model hyper-parameters to gain an estimate of trait heritability. We find that in our variational framework this approach underestimates trait heritability, due to the tendency of mean field variational inference to underestimate the posterior variance of each parameter. Instead we treat the posterior mean 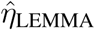 as a fixed effect, and use randomized HE (RHE) regression^23, 24, 64^ to estimate heritability with a single SNP component^23^ (RHE-SC) and multiple SNP components^24^ (RHE-LDMS). When using the multi-component model, SNPs are stratified into a total of 20 bins; using 5 MAF bins (≤ 0.1, 0.1 < MAF ≤ 0.2, 0.2 < MAF ≤ 0.3, 0.3 < MAF ≤ 0.4, 0.4 < MAF ≤ 0.5) and 4 LD score quantiles.

The single component model is given by

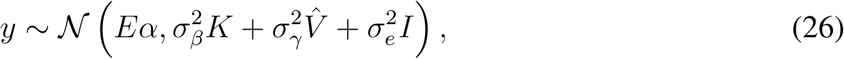

where 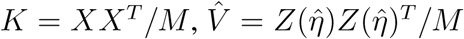 and 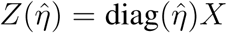. Haseman-Elston (HE) regression is a method of moments estimator that fits the variance components 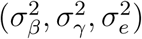 to minimize the difference between the empirical and expected covariances. This is mathematically equivalent to solving the following linear system

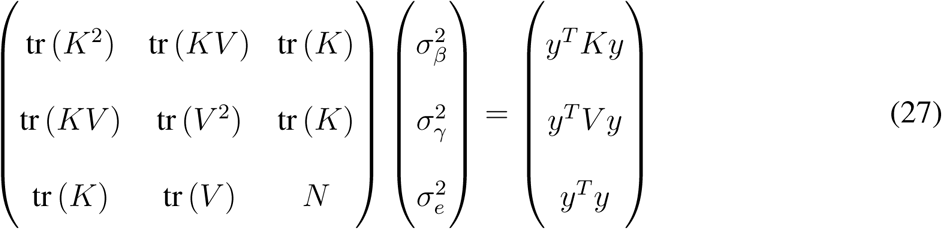

Wu *et al*.^23^ showed that this system can be solved in *𝒪* (*NMB*) time (for small *B*) without ever forming the kinship matrices *K* and *V* using Hutchinson’s estimator, and that covariates can be efficiently projected out of the phenotype, genotypes and interaction matrix *Z* with minimal additional cost. Pazokitoroudi *et al*.^24^ give an extension to multiple components and showed that variance estimates can be obtained with the block jackknife.

Speed *et al*.^27^ show that the usual form for 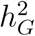, the proportion of trait variance explained by additive genetic effects, given by

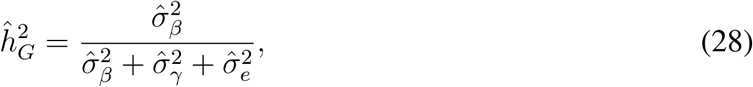

holds only when genotype matrix *X* is standardized to have column mean zero and column variance one. Whilst this is true in expectation for 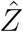 (assuming that 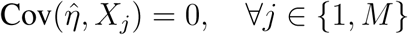), this is not guaranteed. To obtain column mean zero we include an intercept of ones among the covariates that are projected out of the phenotype, genotypes and interaction matrix. To account for columns having variance not equal to one, we use a more general form of the heritability estimator (see Speed *et al*.^27^ for details)

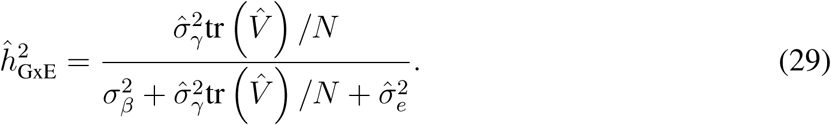

### Implementation and computational efficiency

We provide software implementing the LEMMA algorithm in C++ from https://jmarchini.org/lemma/. We implement a number of steps to improve computational and memory efficiency including vectorization using SIMD extensions, compressed data formats, pre-computing quantities, parallel computing with OpenMPI, use of the well optimized Intel Math Kernel Library. Full details are given in the **Supplementary Material**.

### Detecting squared environmental dependence

By default, each of the *L* environmental variables is tested against the phenotype for significant squared effects. To do this LEMMA tests the hypothesis *β*_*l*_ ≠ 0 using the following linear model

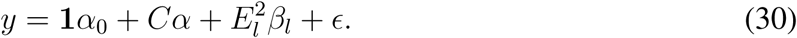

The squared effect of any environmental variables with a p-value less than 0.01 (Bonferroni correction for *L* multiple tests) are added to the matrix of covariates *C*.

### Controlling for covariates

Unlike in BOLT-LMM ^13^, it is not possible to efficiently project covariates out of the model (*y, X, Z*), because the multiplicative interaction matrix *Z* changes after each pass through the data. Instead the LEMMA software package can either regress covariates out of the phenotype or model the covariates as random effects in the variational framework. For our analyses of the UK Biobank we included all covariates within the variational model.

### Comparison to existing GxE methods

We compare LEMMA to three other single SNP methods that jointly model interactions with multiple environments. The first comparison method, StructLMM ^7^, is a method that uses a random effects term *u* to model environmental similarity instead of genetic similarity. Specifically, StructLMM uses the model

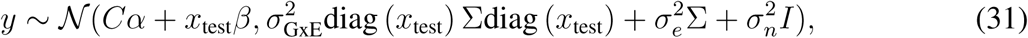

to test the hypothesis 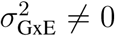. Here *C* is the matrix of covariates with fixed effects *α, x*_test_ is the focal variant and ∑ = *EE*^*T*^ is the environmental similarity matrix (where *E* is an *N* × *L* matrix of environmental variables). Although StructLMM provides both an interaction test and a joint test that looks for non-zero main and interaction effects at each SNP, we use only the interaction test in our comparisons. Finally we note that StructLMM recommends ‘gaussianizing’ the phenotype as a pre-processing step; however we just center and scale the phenotype for consistency with our other methods.

Our second and third comparison methods use equivalent information to StructLMM in a fixed effects framework. Consider the linear model

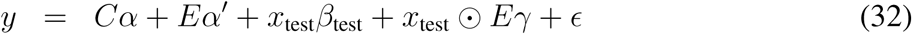

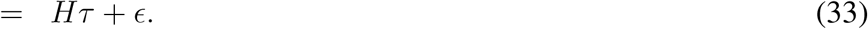

where *H* is formed from column-wise concatenation of [*C, E, x*_test_, diag(*x*_test_)*E*] and *τ* is the corresponding vector of fixed effects. Let *R* be the indicator matrix such that *Rτ* = *γ*. We wish to test the null hypothesis *H*_0_ : *γ* = 0. Assuming that *ϵ* has mean zero and covariance matrix Ω we can use the standard OLS estimator 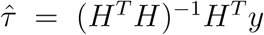 which (under certain regularity conditions) is asymptotically distributed as normal with mean *τ* and variance given by 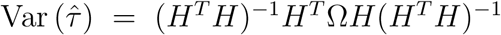. Assuming homoskedasticity yields the standard F test statistic

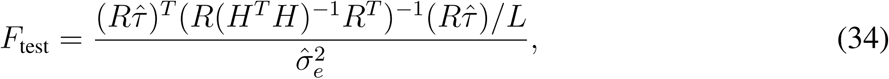

which follows an 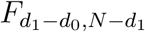 distribution under the null hypothesis, where *d*_1_ is the column rank of *H* and *d*_0_ is the column rank of H under the null hypothesis. Alternatively we can use the same robust standard error used in the LEMMA test statistic

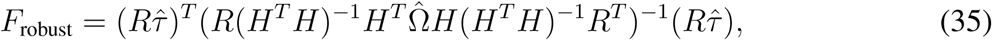

where 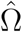 is a diagonal matrix with 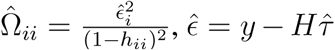 and *h* = *H*(*H*^*T*^ *H*)^−1^*H*. Then *F*_robust_ is asymptotically distributed as 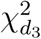 where *d*_3_ is the rank of *HR*^*T*^. In our simulations we refer to this as the robust F-test.

### SNP specific interaction profile

The SNP specific interaction profile is defined as *η*_*LS*_ = *Ew*_*LS*_ where *w*_*LS*_ is the least squares parameter estimate of *w* in the single SNP model

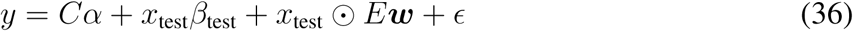

and *y, C* and *E* are the data matrices defined in **Online Methods**. The correlation between *η*_*LS*_ for a given SNP and the ES estimated by LEMMA can be viewed as a proxy for how well LEMMA captures the GxE interactions at that locus.

### UK Biobank analysis

We used real genotype and phenotype data from the UK Biobank, which is a large prospective cohort study of approximately 500, 000 individuals living in the UK ^3^. To account for potential confounding effects of population structure, we first subset down to the white British subset of 344,068 individuals used by Bycroft *et al*.^3^ in a GWAS on human height. This represents unrelated individuals who self-report white British ethnicity and whose genetic data projected onto principal components lies within the white British cluster^3^. After sub-setting down to individuals who had complete data across the phenotype, covariates and environmental factors (see below) we were left with approximately 280, 000 samples per trait (**Table S1**). Finally we filtered genetic data based on minor allele frequency (≥ 0.01) and IMPUTE info score (≥ 0.3), leaving approximately 642, 000 genotyped variants (**Table S1**) and 10, 295, 038 imputed variants per trait. For each trait we included age^3^, age^2^ × gender, age^3^ × gender, a binary indicator for the genotype chip and the top 20 genetic principal components as additional covariates.

BMI was derived from height and weight measurements made during the first assessment visit (instance ‘0’ of field 21001), and readings more than six standard deviations from the population mean to missing. logBMI and INT(BMI) refer to BMI after applying a log transformation and an inverse normal transformation (applied separately to males and females) respectively.

After calculating the mean SBP and DBP using automated blood pressure readings from the first assessment visit (fields 4080 and 4079), we adjusted for medication usage by adding 15mmHg and 10mmHg to SBP and DBP respectively ^65^. Data from manual measurements (fields 93 and 94) was used in the rare instance that no automated reading was available. Blood pressure readings more than four standard deviations from the mean were set to missing. PP was then calculated as SBP minus DBP.

For our GxE analyses, we made use of 42 environmental variables from the UK Biobank, similar to those used in previous GxE analyses of BMI in the UK Biobank ^7, 26^. From the data provided by the UK Biobank we included 7 continuous environmental variables (“Age when attended assessment centre”, “Sleep duration”, “Time spent watching television”, “Number of days/week walked 10+ minutes”, “Number of days/week of moderate physical activity 10+ minutes”, “Number of days/week of vigorous physical activity 10+ minutes”, “Townsend deprivation index at recruitment”), 1 ordinal environmental variable (“Alcohol intake frequency”), 9 dietary ordinal variables (“Salt added to food”, “Oily fish intake”, “Non-oily fish intake”, “Processed meat intake”, “Poultry intake”, “Beef intake”, “Lamb intake”, “Pork intake”, “Cheese intake”) and 2 dietary continuous variables (“Tea intake”, “Cooked vegetable intake”). We further derived 1 categorical variable (“Is Current Smoker” from the responses given in the UK Biobank field “Smoking status”) and 1 continuous variable (“Sleep sd”; the number of standard deviations from the population mean sleep duration). For analyses of blood pressure, we additionally included one further continuous variable “Waist circumference”. This left 11 dietary variables and 10 non-dietary variables (11 for blood pressure traits). In addition we included multiplicative interactions between participants age and gender with all non-dietary variables, and included the main effect of gender, giving the data matrix *E* a total of 42 columns (45 for blood pressure traits). Before running LEMMA each column was standardized as

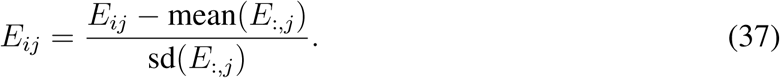

In all cases, where participants responded with “Prefer not to answer”, “Do not know” or “None of the above” we set the value to missing. For 3 continuous variables (“Time spent watching television”, “Tea intake”, “Cooked vegetable intake”) we removed the 99’th percentile and for “Sleep duration” we removed both the 1st and 99th percentiles.

After running LEMMA, we found it convenient to interpret weights corresponding to a rescaled data matrix *E*_1_. Assuming the column space of *E* and *E*_1_ is the same, weights ***w***_**1**_ that correspond to *E*_1_ can be extracted from the ES using least squares

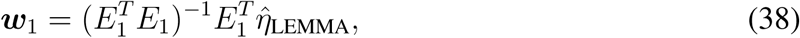

where 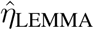 represents the ES. We note that although multivariate linear regression is invariant to a re-scaling of the design matrix, ridge regression is not due to the penalization place on the magnitude of the learned parameters. However, as the magnitude of the weights from our UK Biobank analysis is typically small (less than 0.2) compared to the standard deviation of our Gaussian prior (1) in this case the re-scaling makes minimal difference.

Re-coded data matrix *E*_1_ was formed with one column for each of the 11 dietary variables (normalized to have mean zero and variance one), and three columns for each of the 10 (11 for blood pressure traits) non-dietary variables; the first augmented by a binary male indicator vector, the second by a binary female indicator vector and the third by a continuous vector of participant age. Columns augmented by male and female binary indicator vectors were normalized to have mean zero and variance one (not including zeros due to augmentation), apart from age (scaled to represent the number of decades aged past 40). Columns augmented by age were normalized first and then multiplied by age on the per decade scale. We further included indicator columns for men and women, which can be interpreted as gender specific intercepts and is equivalent to including an intercept and a binary column for only one gender (men or women). We note this leaves 43 (46) columns where the extra column comes from including an intercept within the column space of *E*_1_ and is necessary because some columns have mean not equal to zero. Thus the column space of *E*_1_ is equivalent to *E* under the constraint that the ES has mean zero.

### Simulation studies

Genetic data was sub-sampled from the UK Biobank, by default using *N* = 25*K* unrelated individuals of mixed ancestry and *M* = 100*K* genotyped SNPs. Environmental variables were simulated from a standard gaussian distribution. By default we constructed phenotypes with 2500 causal main effects and 1250 causal interaction effects explaining 20% and 5% of trait variance respectively. For each phenotype we constructed a weighted average of the environmental variables which we used to simulate multiplicative interaction effects. Environments with a non-zero weight are referred to as active. All non-zero effects were drawn from SNPs in the first half of each chromosome, allowing us to test the calibration of each method on ‘null’ SNPs from the second half of each chromosome. To allow direct power comparisons across different scenarios we included an additional 60 SNPs with standardized effect sizes, that together accounted for 1% of trait variance with their main effects and 1% of trait variance with their interaction effects. Finally, a further 1% of trait variance was modeled using the first genetic principal component (PC). For all methods we included the first genetic PC as a covariate. For each method we calculated power as the proportion of the SNPs of standardized effect identified at a threshold of *p* < 0.01.

In simulations used to test randomized HE-regression, phenotypes were constructed with 10, 000 causal main effects explaining 20% of trait variance and, in simulations with non-zero GxE heritability, 10, 000 causal SNPs with interaction effects.

### Model misspecification

We simulated a scenario where a disease trait *Y* depends non-linearly on a heritable environmental factor *S*. More explicitly suppose that *X* is the centered and scaled genotype matrix so that columns have mean zero and variance one, that *S* is modeled as

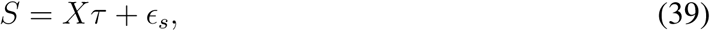

where 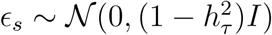, *τ* models random SNP effects for *S* and trait *Y* is given by

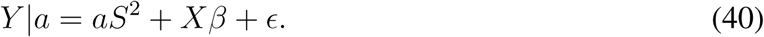

Here *a* is a constant that we use to control the strength of the contribution of *S*^2^ to *Y*, 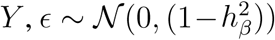 and *β* is the random SNP effects for *Y*. For simulation we suppose that *τ* and *β* have spike and slab priors

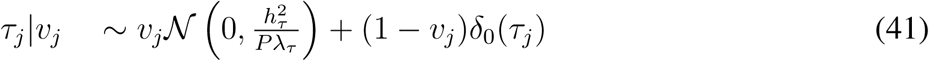

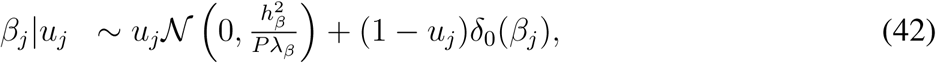

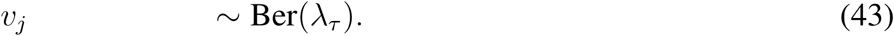

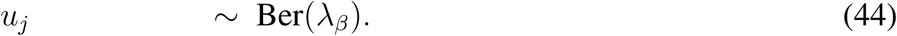

### Data availability

The genetic and phenotype datasets generated by UK Biobank analyzed during the current study are available via the UK Biobank data access process. The Resource is available to all bona fide researchers, from academic, charity, public, and commercial sectors, for all types of health-related research that is in the public interest, without preferential or exclusive access for any person. More details are available here http://www.ukbiobank.ac.uk/register-apply/

## URLs

UK Biobank: http://www.ukbiobank.ac.uk

LEMMA: https://jmarchini.org/lemma/

StructLMM as implemented in LIMIX 2.0.0: https://github.com/limix/limix

## Acknowledgments

Computation used the Oxford Biomedical Research Computing (BMRC) facility, a joint development between the Wellcome Centre for Human Genetics and the Big Data Institute supported by Health Data Research UK and the NIHR Oxford Biomedical Research Centre. Financial support was provided by the Wellcome Trust Core Award Grant Number 203141/Z/16/Z. The views expressed are those of the author(s) and not necessarily those of the NHS, the NIHR or the Department of Health. We are grateful to Kevin Sharp, David Steinsaltz and Helen Warren for discussions about this work.

## Author contributions

J.M. and M.K. conceived the ideas for the model and methods development. M.K. conducted all analyses, derived the variational Bayes updates and developed the software that implemented the methods with guidance from J.M. M.K. and J.M. wrote the manuscript. J.M. carried out this work while affiliated with the University of Oxford.

## Supplementary Material

### Derivation of Variational Bayes updates

We use Coordinate Ascent Variational Inference (CAVI) to optimize the ELBO^66^. CAVI is a cyclic optimization strategy that iteratively maximizes the ELBO with respect to each latent variable whilst holding the others fixed. We now provide a brief justification of the CAVI update step and then derive the update for each of the latent variables in the LEMMA model.

Using the fact that the variational distributions factorizes, we can write the ELBO as

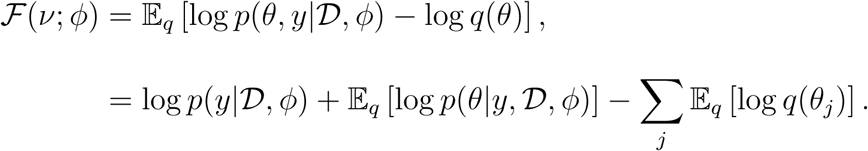

Hence it is relatively simple to extract out dependance of ℱ(*ν*; *ϕ*) on *θ*_*j*_

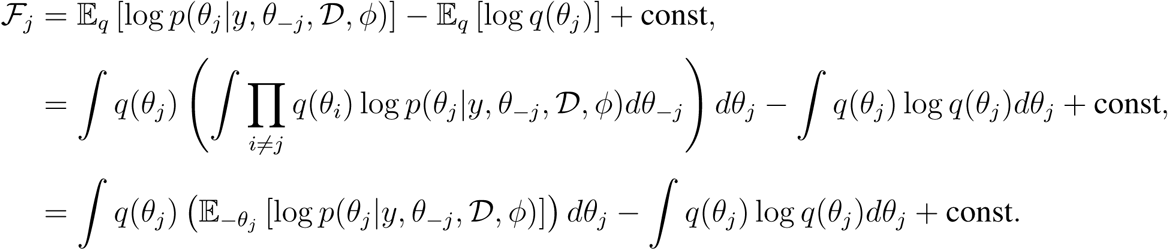

The last line is proportional to the KL divergence between log *q*(*θ*_*j*_) and 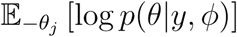, where 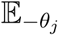 denotes the expectation with respect to the *q* distributions over all variables {*θ*_*i*_ : *θ*_*i*_ ≠ *θ*_*j*_}. Therefore to maximize the ELBO with respect to *q*(*θ*_*j*_) we must minimize the KL divergence between log *q*(*θ*_*j*_) and 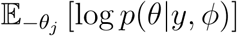, which occurs when

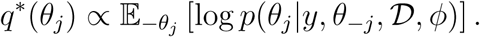

After applying Bayes theorem, the above CAVI step can be equivalently expressed as

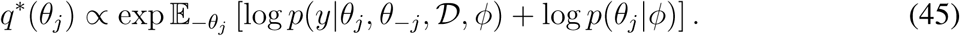

### Updates for SNP main effect sizes *q*(*β*_*j*_)

The prior and conditional log-likelihood for *β*_*j*_ are given by

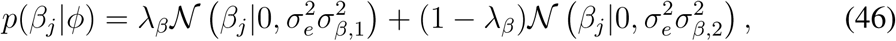

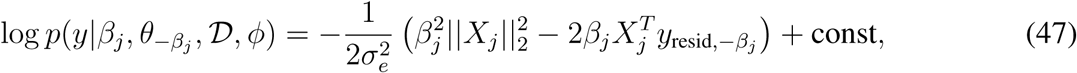

where const is a constant independent of *β*_*j*_ and 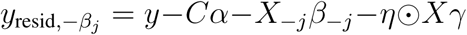. Substituting eq. (47) and eq. (46) into eq. (45) yields

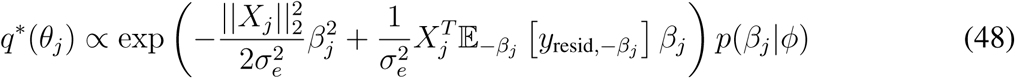

as the prior is independent of 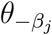. We now note the result

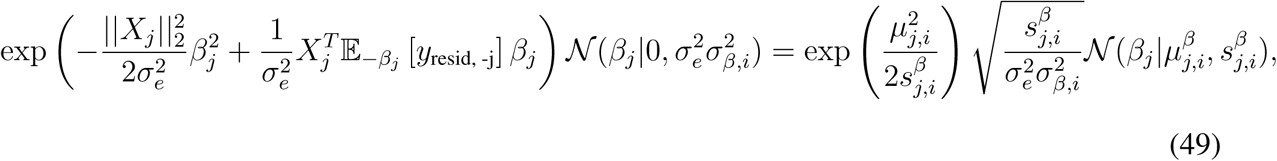

where

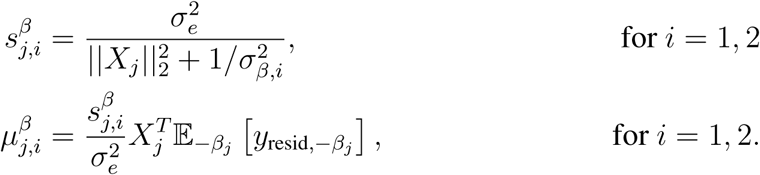

Substituting eq. (49) into eq. (48) yields

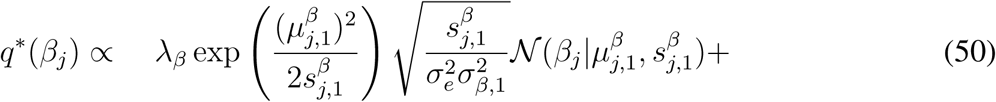

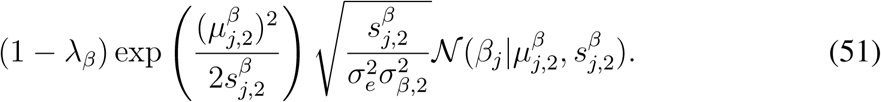

It is now clear that *q*^∗^(*β*_*j*_) is the probability density function of a mixture of gaussians

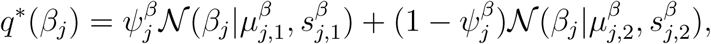

where the mixture components 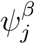 and 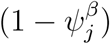 must sum to one. Therefore

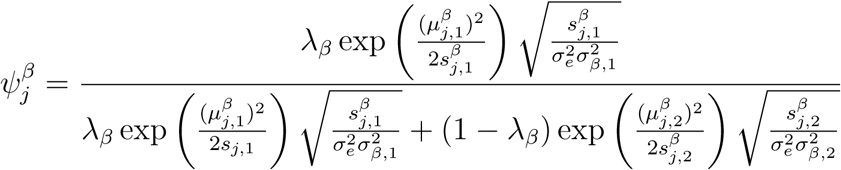

or equivalently

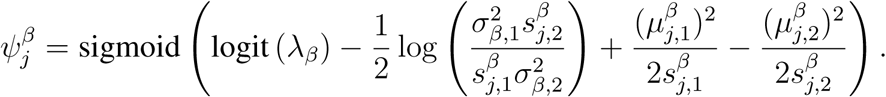

Therefore, the update equations for *q*(*β*_*j*_) can be summarised as

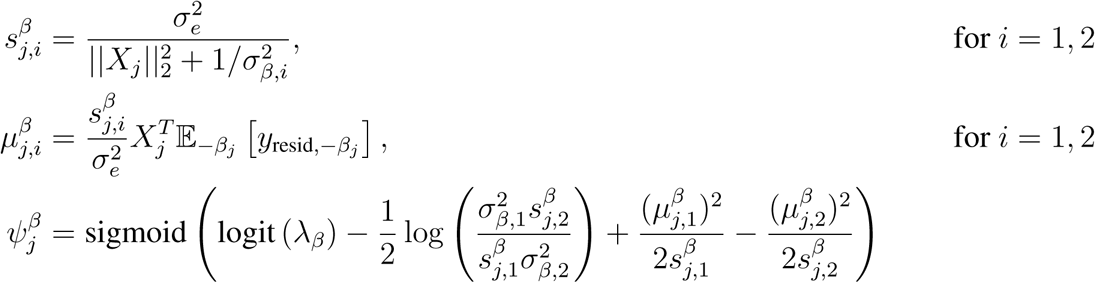

where

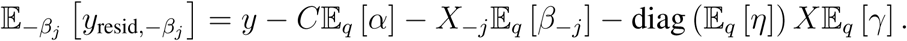

### Updates for SNP interaction effect sizes *q*(*γ*_*j*_)

The derivation of the variational update for *q*(*γ*_*j*_) is extremely similar to that of *q*(*β*_*j*_). The prior and conditional log-likelihood for *γ*_*j*_ are given by

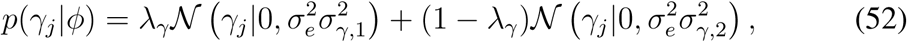

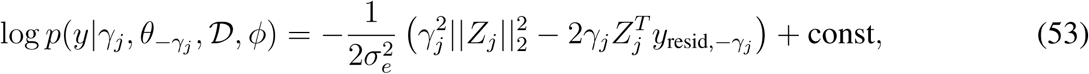

where const is a constant independent of *γ*_*j*_ and 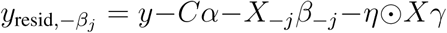. Substituting eq. (53) and eq. (52) into eq. (45) yields

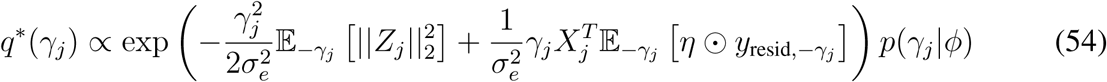

as the prior is independent of *θ*_−*j*_. Following the same steps as used in the derivation of *q*^∗^(*β*_*j*_), is it clear that *q*^∗^(*γ*_*j*_) is also the probability density function of a mixture of gaussians

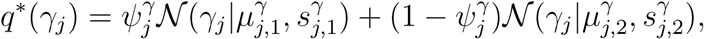

whose optimal CAVI updates are given by

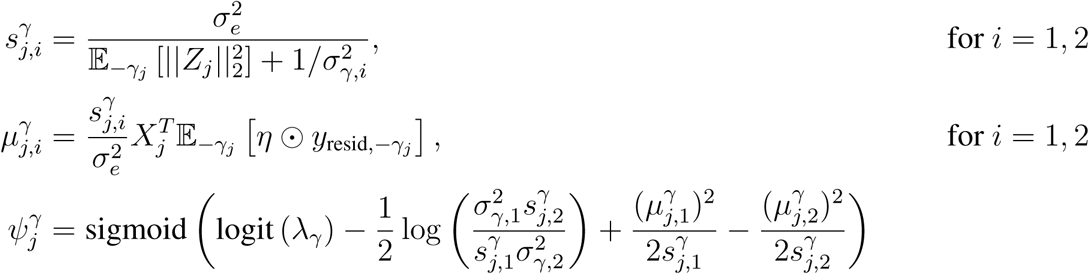

where

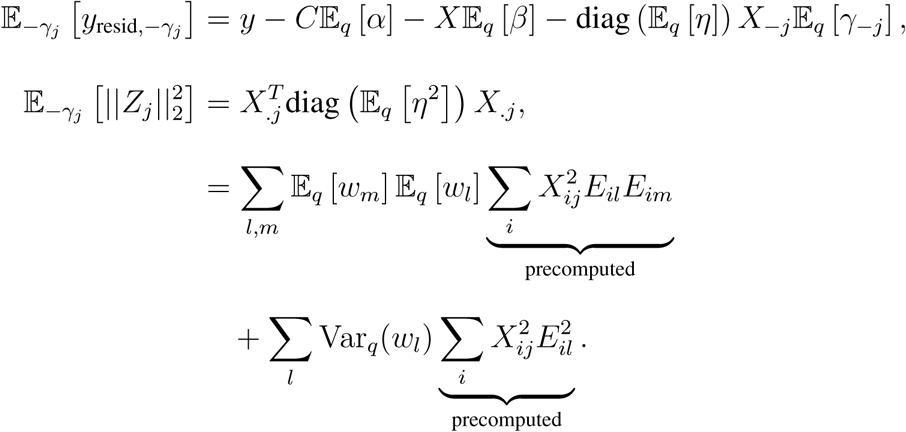

Note that computation of 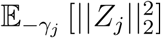 is an *O*(*L*^2^ + *N*) operation due to the precomputation of 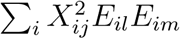 (and without this precomputation the compute cost of this update would be 𝒪(*NL*^2^)).

### Updates for interaction weights *q*(*w*_*l*_)

Rewriting the conditional log-likelihood makes its dependence on *w* clear

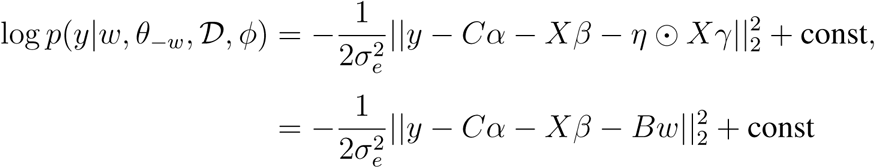

where *B* = diag (*Xγ*) *E* and const is a constant independent of *w*. For convenience we denote the *l*’th column of *B* as *B*_*l*_. Therefore the prior and conditional log-likelihood of *w*_*l*_ are given by

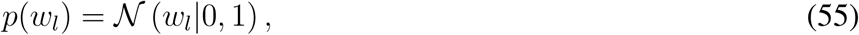

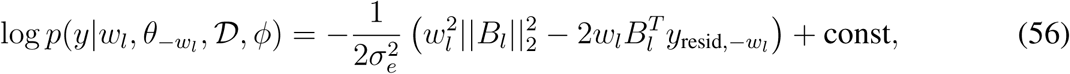

where 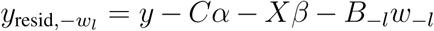 and const is now a constant independent of *w*_*l*_. Substituting eq. (56) and eq. (55) into eq. (45) yields

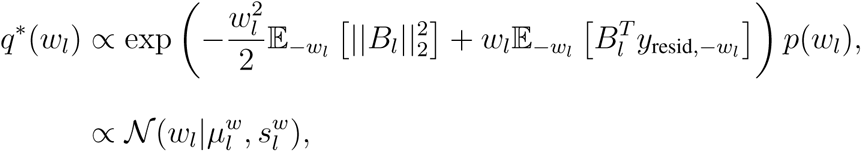

where

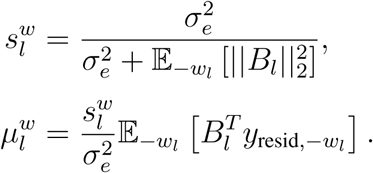

As *q*^∗^(*w*_*l*_) must be a valid distribution, it is clear that 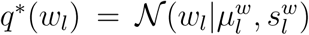. The quantities 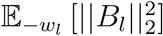 and 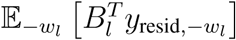 can be computed as follows

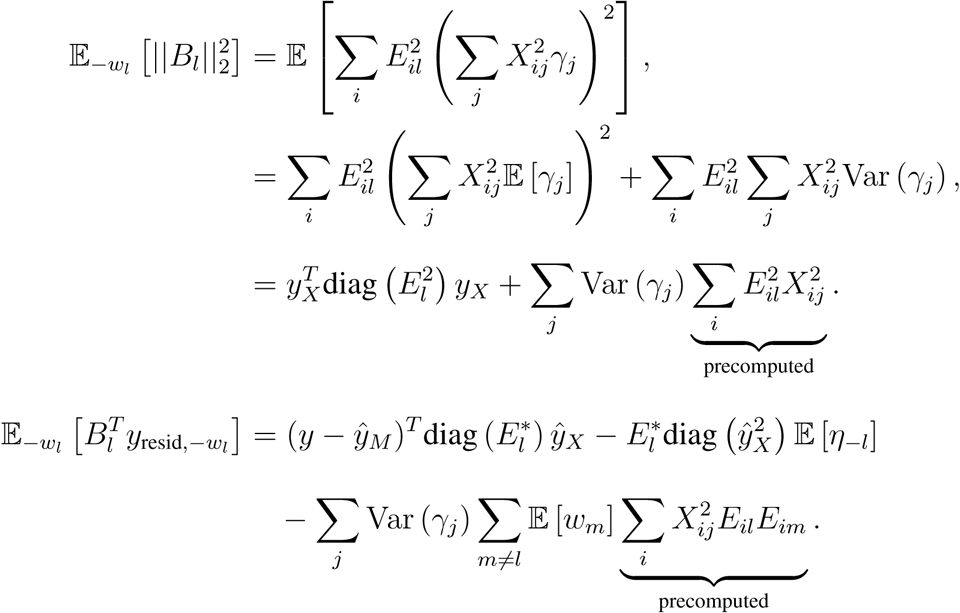

Note that computation of 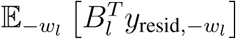 is an *O*(*NL*) operation due to the precomputation of 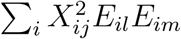 (and without this precomputation the compute cost of this update would be 𝒪(*NML*)).

### Updates for covariate main effect sizes *q*(*α*_*c*_)

The derivation of the variational update for *q*(*α*_*c*_) is extremely similar to that of *q*(*w*_*l*_). The prior and conditional log-likelihood for *α*_*c*_ are given by

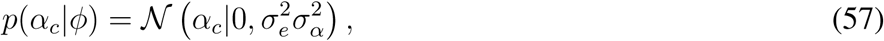

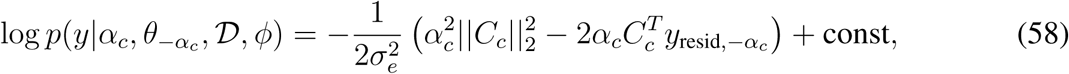

where const is a constant independent of *α*_*c*_ and 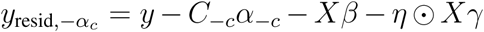. By similarity with the derivation of *q*^∗^(*w*_*l*_) it is clear that *q*^∗^(*α*_*c*_) is a gassian distribution, with variational updates

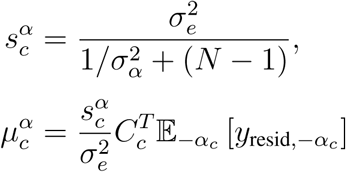

where 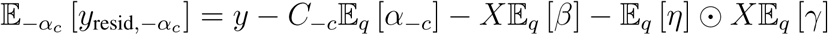.

#### Evidence lower bound

Variational inference involves maximising the evidence lower bound (ELBO) ℱ(*ϕ*; *ν*) on the model log-likelihood log *p*(*y*|𝒟, *ϕ*). The ELBO can be separated into the expected conditional log-likelihood and the KL divergence between the variational distribution and the respective priors. This is given by

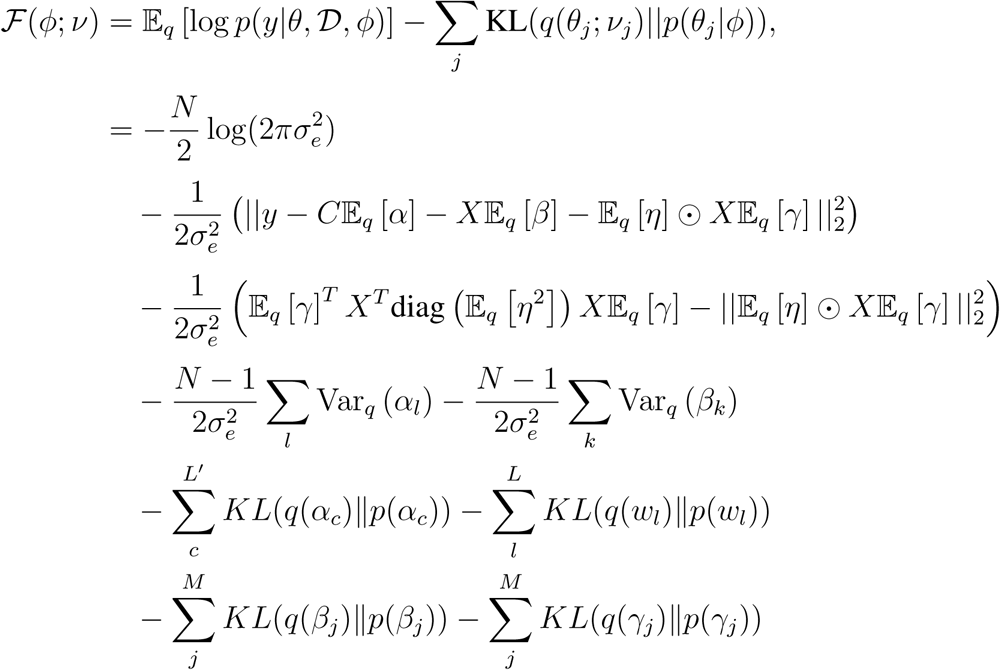

While the KL Divergence between two univariate gaussian distributions is a standard result, the KL Divergence between two mixtures of gaussians is not analytically tractable. However the matched bound approximation ^67^ can be used to provide an upper bound when both have the same number of components. Thus for two mixtures of gaussians given by

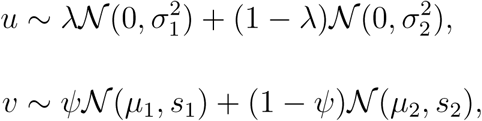

the matched bound on the KL divergence is given by

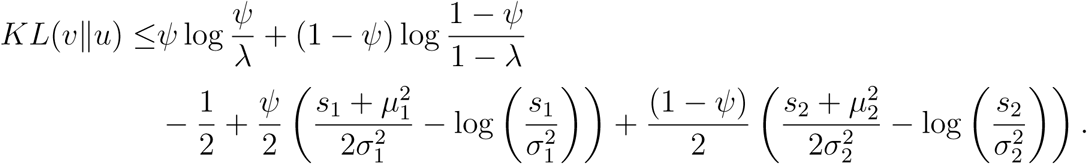

Use of the matched bound approximation retains a valid variational algorithm, because it maintains the lower bound on the marginal log-likelihood^68^.

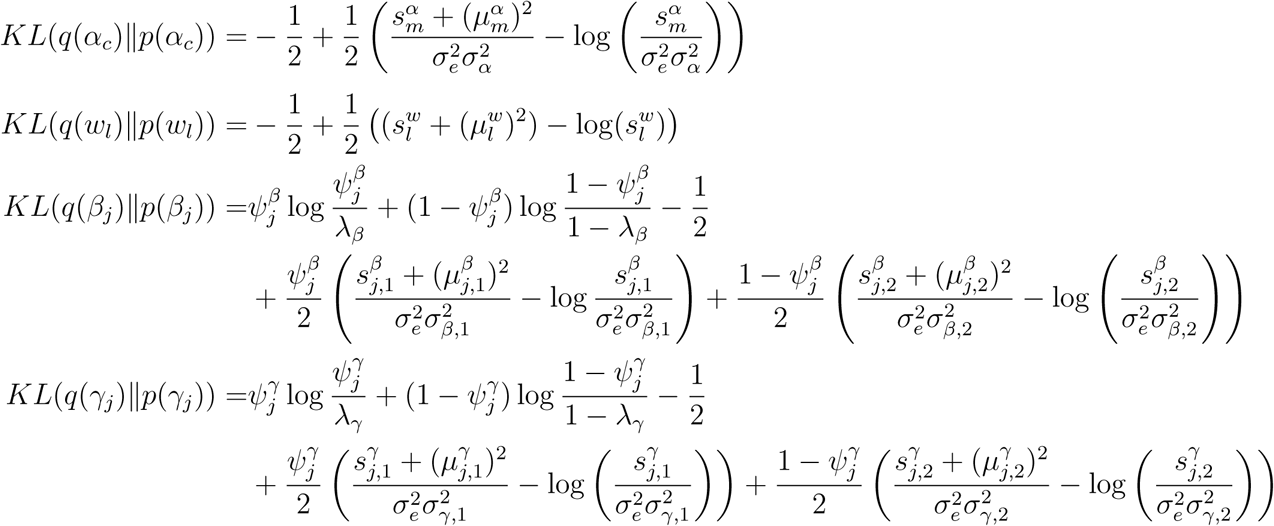

#### Derivation of hyperparameter maximization

For the maximization step we set 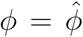 where ∇_*ϕ*_*F* (*ϕ*; *ν*) = 0. For ease of notation we perform the following change of variables

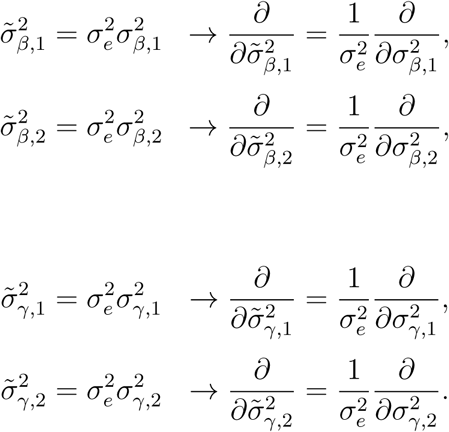

This makes the derivation easier as all the partial derivatives become decoupled. Partial derivatives with respect to each hyper-parameter are given by

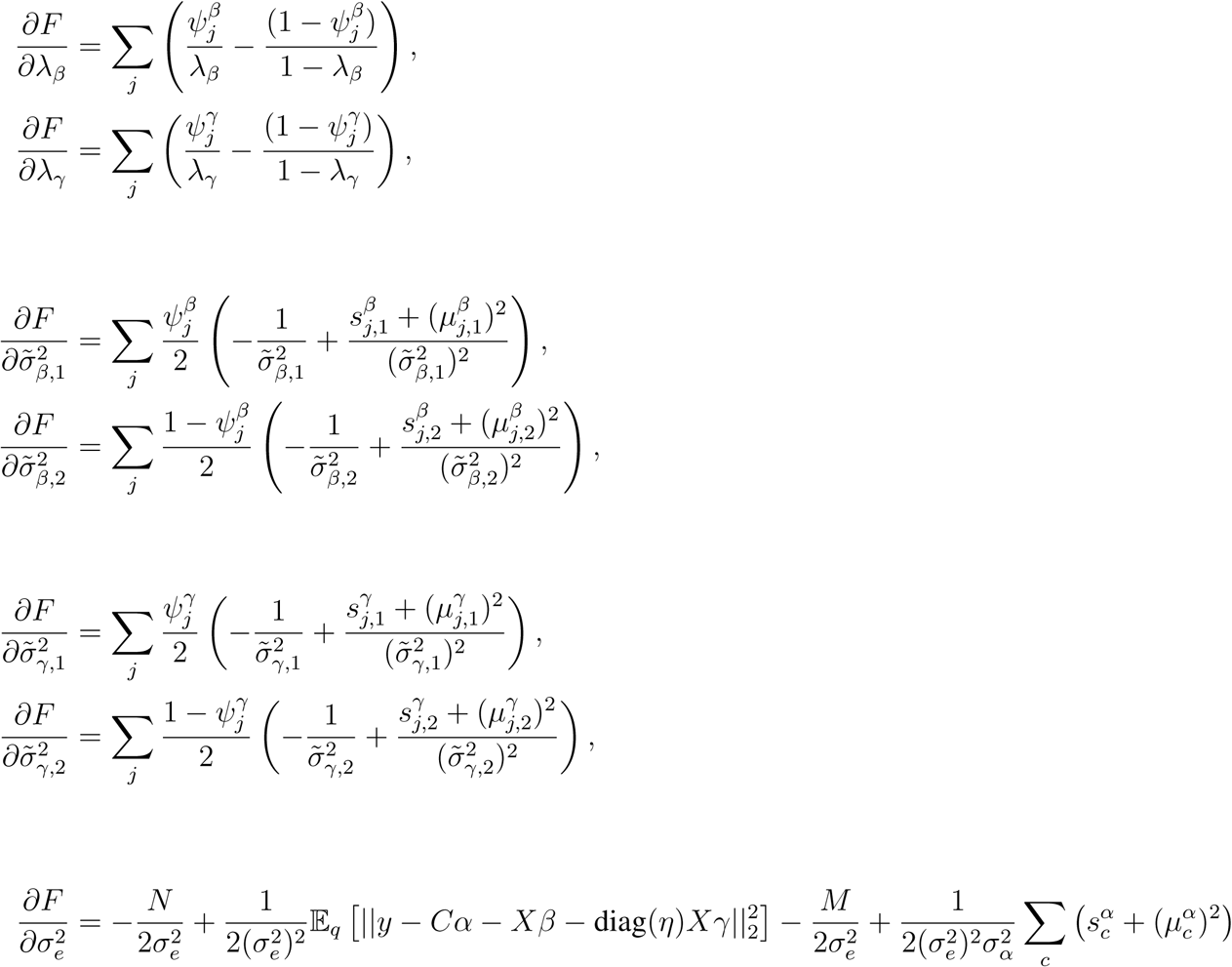

Hence the maximization steps are

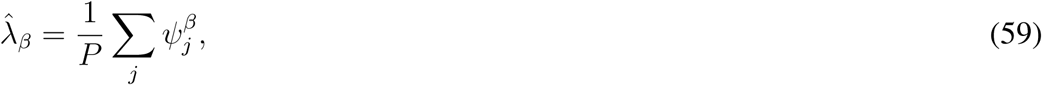

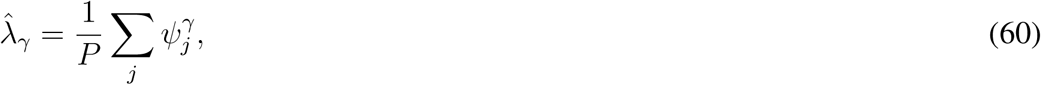

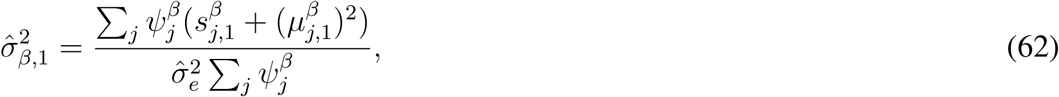

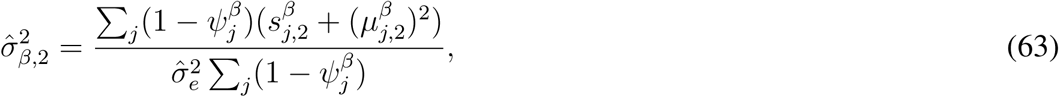

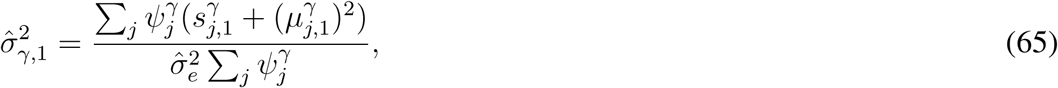

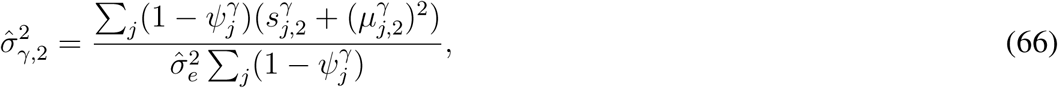

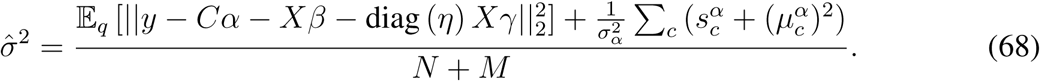

As an aside we note that one could use the maximized hyper-parameters (after convergence) to obtain a point estimate of Var (*β*). However, by substituting in Equations (59) to (66) we can see that this is equivalent to 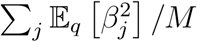.

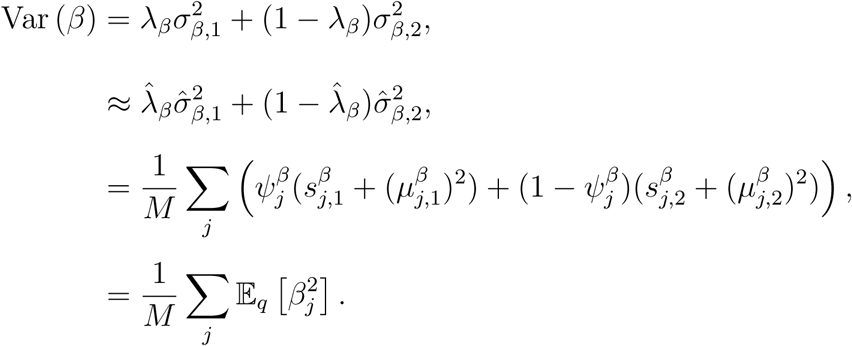

As the mean field assumption tends to cause variational inference algorithms to underestimate the variance of latent variables ^69^, this is likely to produce an underestimate of Var (*β*). We can observe the same result for Var (*γ*) with an analogous argument.

#### Compressed genotype data

To reduce RAM usage, LEMMA stores a compressed version of the genotype matrix using *NM* bytes. To do this LEMMA splits the interval [0, 2] into 2^8^ segments and stores the index of the segment that each dosage falls into, as well as the mean and variance for each SNP. Then when operating on a SNP, LEMMA reconstructs the centered and scaled dosages for that SNP. This approach is similar to that used by the BGEN data format ^70^ and results in a small loss of accuracy, but is more flexible that assuming dosages are hardcoded to {0, 1, 2}.

#### Computational efficiency

Using mean field variational inference, estimation of the posterior means of the latent variables *β, γ, w* can be reduced to an iterative algorithm that cycles through the variables sequentially, updating each conditional on the values of the others. Taking the main effect of the *j*’th SNP as an example, the update scheme for *β*_*j*_ can be written heuristically as

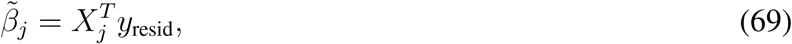

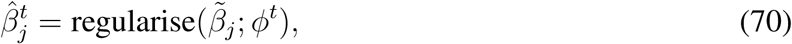

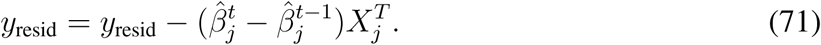

In Equation (69) we compute the correlation between the *j* SNP and the residual phenotype vector. In Equation (70) we compute the posterior mean of *β*_*j*_ which depends on the correlation with the residual phenotype, the prior on *β*_*j*_ and the current hyper-parameters Finally in Equation (71) we update the residual phenotype vector.

The majority of computational time is spent on the dot product in Equation (69) and updating the residual phenotype in Equation (71). Both are BLAS Level 1 operations, which implies that memory access is often the principle bottleneck rather than the number of cores available. It is possible to step up to BLAS Level 2 by updating a block of SNPs in parallel ^13^, however this is still a memory bound operation. Instead we use a parallel computing strategy suggested by ^71^ for use in genome wide regression, and subsequently used by ^32^, to compute the dot product and perform the residual update in parallel using OpenMPI. Briefly, we partition the samples such that blocks of rows of the phenotype *y*, genotypes *X* and environmental variables *E* are assigned to each core. For a given update step, each core calculates the dot product for the locally held block of samples and then shares the local dot product with the rest of the network. From this the dot product for the entire cohort can be reconstructed cheaply. After computing the posterior mean, each core then updates the residual phenotype for the block of samples stored locally. We observed that a distributed algorithm using OpenMPI was faster than the same algorithm using multi-threaded matrix-vector operations with the Intel MKL Library even on a single node with multiple cores. However using OpenMPI has the additional advantage of allowing users to utilize cores from across a cluster rather than being restricted to a single node. **Figure S18** shows LEMMA scales with increasing sample size.

#### Pre-computed quantities

To aid computational efficiency we pre-compute a *M* ×*L*(*L*+1) matrix *W* where

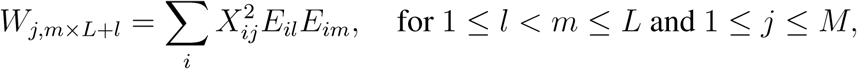

and is used in the updates of *q*(*w*_*l*_) and *q*(*γ*_*j*_). LEMMA can compute this internally, incurring a one off cost of 𝒪(*NML*^2^), or is able to read from a text file at run time. As this is easily computed in parallel over batches of variants and/or environment, we recommend that for biobank scale datasets users should pre-compute this quantity beforehand using a separate tool that we have provided.

#### Parameter Initialization

We start the variational mean estimates of *q*(*β*) and *q*(*γ*) at zero. To initialize mean estimates of the interaction weights *q*(*w*) we have two options; the first of which is simply to use a uniform weighting over all environments. For the second we apply an F-Test independently at each SNP and use the learned coefficients from the test with the lowest p-value as the initial values of the interaction weights. We find that we often obtain similar results from both options, so for simplicity we use a uniform start point for our Biobank analyses. To initialize mean estimates of *q*(*α*) we use the least squares fit of *C* on *y*.

Inintial values of the hyperparameters are drawn randomly from the following distributions

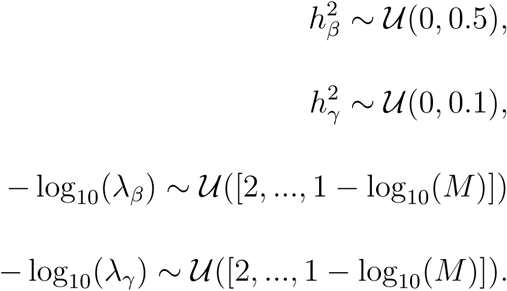

We then set

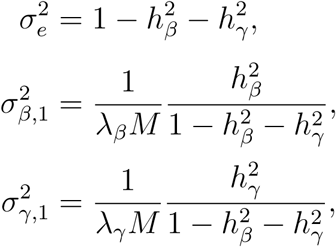

and initialize the spike variances at

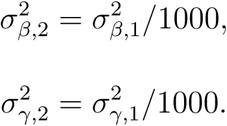

Setting the sparsity hyperparameters *λ*_*β*_, *λ*_*γ*_ in this manner allows LEMMA to start from a state where only a small number (somewhere between ten and one in one hundred) SNPs are expected to be part of the slab prior. The sparsity hyperparameters can then be updated in the variational maximization step to better reflect trait genetic architecture.

#### Missing data

Samples with missing data in the phenotype, environmental variables or covariates are excluded. By default LEMMA imputes missing genetic data with the mean dosage of each SNP, however as LEMMA does not assume dosages are hard called with {0, 1, 2} we recommend that users first impute genetic data with standard imputation pipelines.

#### Robust standard errors in GxE Studies

In **Figure S19** we illustrate how a multiplicative GxE interaction effect on a quantitative trait can cause the conditional trait variance given an interacting SNP Var(*Y* |*g*_0_) to differ according to the interacting SNPs genotype. This is known as conditional heteroskedasticity and is the key insight behind several recent methods to detect SNPs with non-zero GxE effects in the UK Biobank ^8, 40^.

In the same figure, we can observe that the conditional trait variance given the environmental exposure Var(*y*|*E*) also displays signs of conditional heteroskedasticity. Previous studies ^49^ have observed that methods that assume heteroskedasticity can display substantial inflation when testing for GxE effects at SNPs where there is no true GxE effect. In our simulations we observed that inflation of GxE tests statistics from LEMMA-S and the F-test, both of which assume homoskedasticity, increased with SNP-GxE heritability. Below we give an explanation for this phenomenon.

Consider a polygenic quantitative trait *Y* that has multiplicative GxE interactions with the same environmental exposure *E* at multiple SNPs

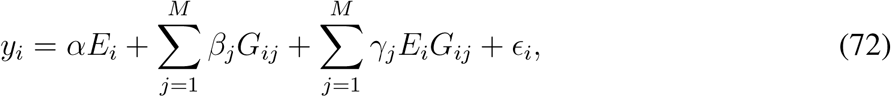

where *M* is the number of SNPs and the coefficients represent true effects. For simplicity we assume that *E* and SNPs *G*_*j*_ are normalized to have mean zero and variance one, that the set of *E* with all causal SNPs {*E*} ∪ {*G*_*j*_ : *β*_*j*_ ≠ 0} is pairwise independent and that the influence from population structure is negligible.

Suppose we have identified *E* as an environmental variable that may plausible have GxE interactions with our phenotype and we then conduct a GWAS for GxE effects. Then at the *k*’th SNP we wish to test the hypothesis *γ*_*k*_ ≠ 0 in the following linear model

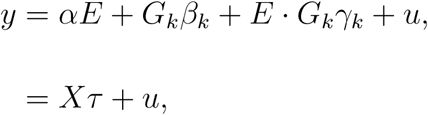

where in the second line *τ* = (*α, β*_*k*_, *γ*_*k*_)^*T*^, *X* is the corresponding design matrix encapsulating all fixed effects and *u* in an unobserved random effects capturing residual noise. Assuming that 𝔼 [*u*|*X*] = 0, the usual least squares estimate of 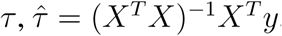, has asymptotic distribution

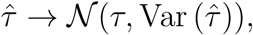

where

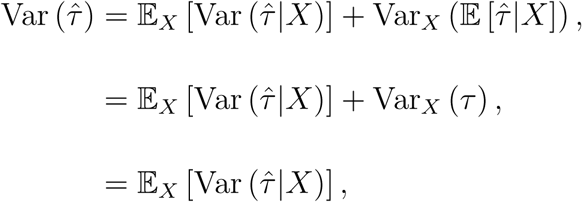

and

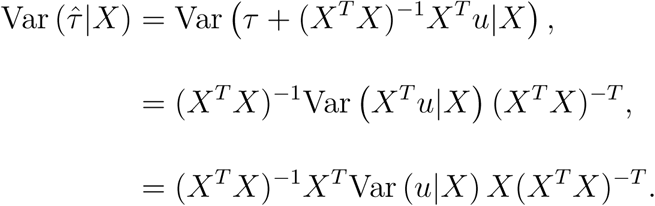

The usual approach is to assume that Var (*u*|*X*) = *σ*^2^*I* (ie homoskedasticity), which yields the standard variance estimator 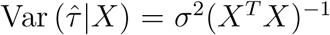. However, given the true generative model for *y* given in Equation (72), we can write *u* as

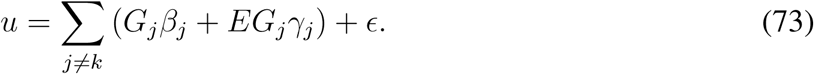

Therefore the conditional variance of *u* given *X* is given by

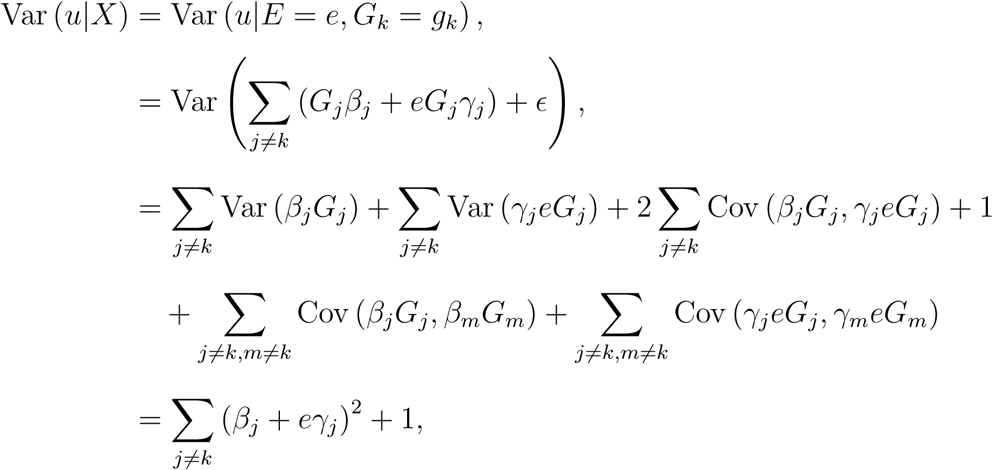

where the covariances in the second line are all zero due to pairwise independence of the set {*E*} ∪ {*G*_*j*_ : *β*_*j*_ ≠ 0}. Thus the conditional trait variance will vary depending on the strength of environmental exposure either if there are a few SNPs with GxE interactions of large effect or if there are many SNPs with small yet non-zero interaction effects, and in either case homoskedasticity is unlikely to be an appropriate assumption.

Robust standard errors, alternatively called Huber-White, sandwich or “heteroskedastic-consistent” errors ^59, 60^, are standard tools used in economics ^47^ to overcome this issue and have previously been proposed for use in GxE interaction studies ^25, 49, 61^. We further include a small adjustment that reduces bias in small samples ^62^. This yields the variance estimator

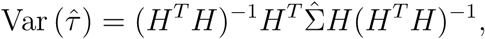

where 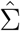 is a diagonal matrix with 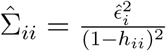, where 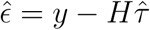 and *h* = *H*(*H*^*T*^ *H*)^−1^*H*^*T*^.

## Supplementary Figures

**Figure S1:**
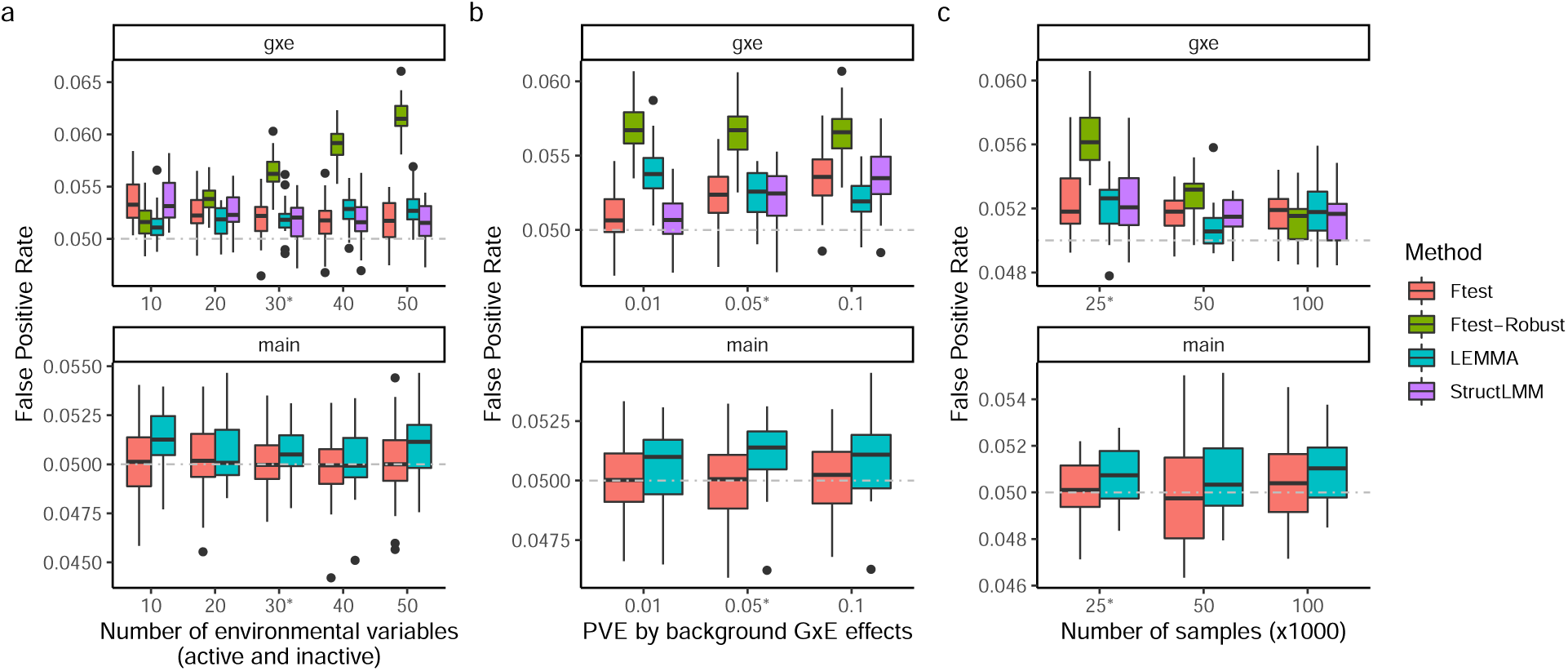
False positive rates on simulated datasets. False positive rate (FPR) for SNP main effects tests (bottom) and SNP GxE interaction tests (top) at null SNPs in the second half of each chromosome, whilst varying (a) the number of environmental variables, (b) proportion of trait variance explained by background GxE effects and (c) sample size. The grey line denotes expected FPR. Simulations used genotypes sub-sampled from the UK Biobank and by default contained *N* = 25*K* samples, *M* = 100*K* SNPs, 6 environmental variables that contributed to the ES and 24 that did not (default parameters denoted by stars). We performed 20 repeats for each scenario. See **Online methods** for full details of phenotype construction.

**Figure S2:**
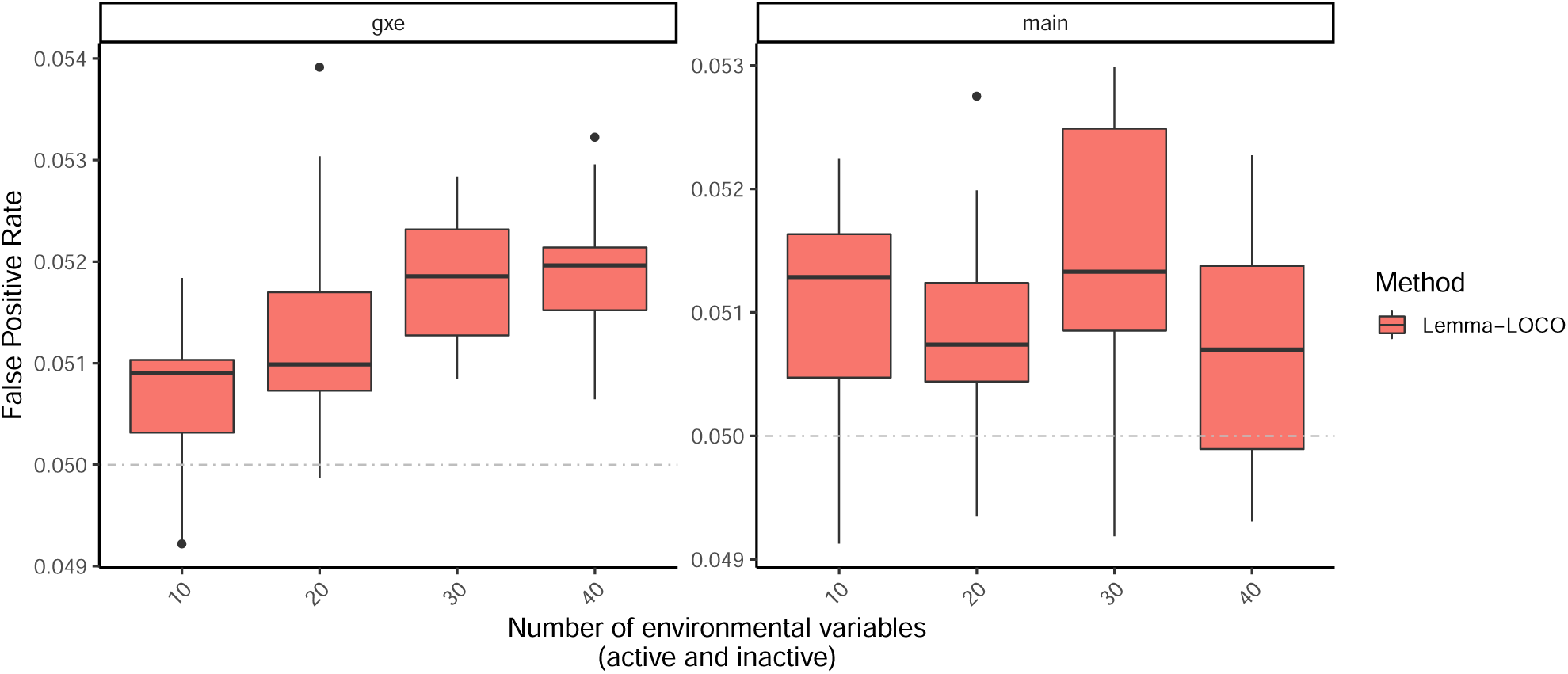
LEMMA false positive rate in large simulations. False positive rate (FPR) for SNP main effects tests (right) and SNP GxE interaction tests (left) at null SNPs in the second half of each chromosome, whilst varying the number of environmental variables. The simulation was conducted with *N* = 200*K* samples and *M* = 400*K* SNPs. The simulated trait was constructed with 10, 000 causal SNPs main effects that explained 20% of variance, and zero causal SNP GxE effects. We performed 20 repeats in each scenario. See **Online methods** for full details of phenotype construction.

**Figure S3:**
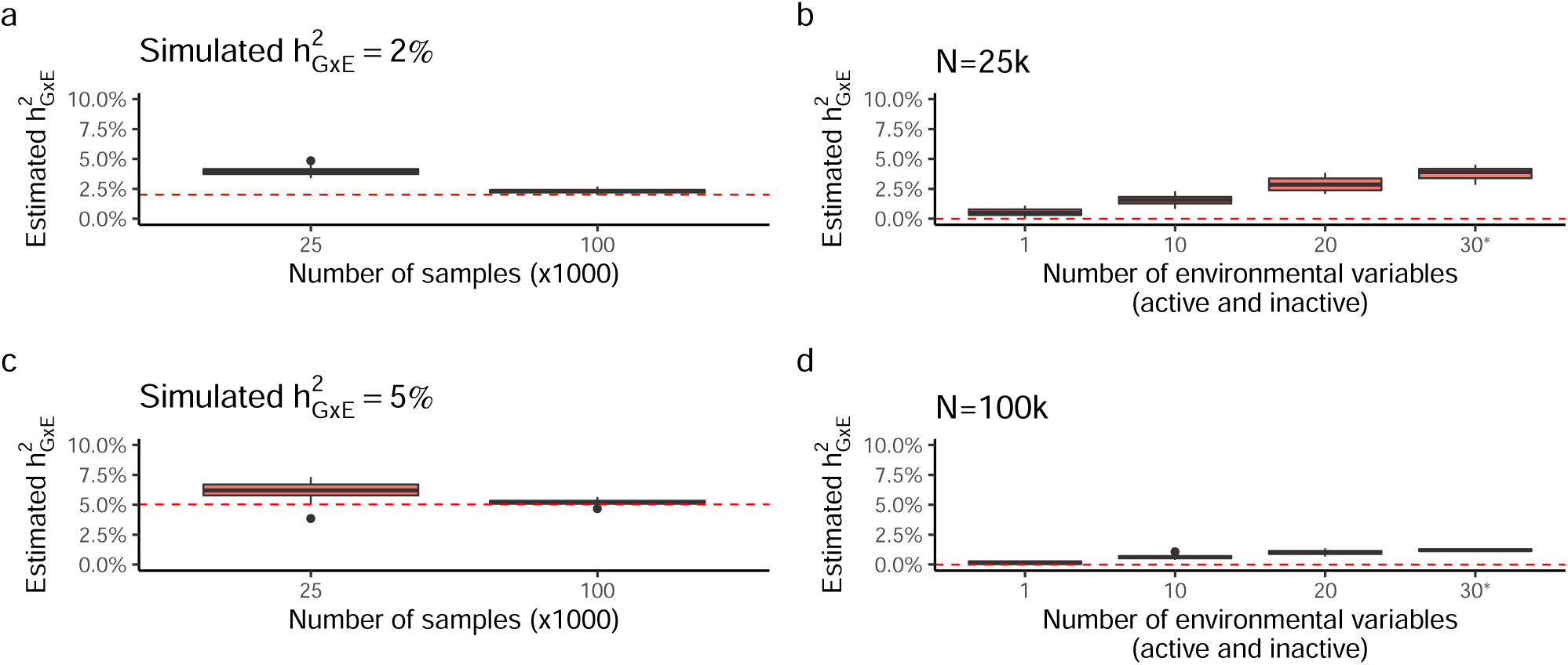
Estimation of GxE heritability. Estimates of SNP-GxE heritability whilst varying the number of environmental variables (b, d) and sample size (a, c). The red dotted line denotes the true SNP-GxE heritability used whilst constructing the simulation. We observed some upwards bias as the number of environmental variables increases (b, d), which is ameliorated with increased sample size (d). Phenotypes were constructed using *M* = 100, 000 SNPs with *M*_causal, main-effects_ = 80, 000 causal main effects and *M*_causal, GxE-effects_ = 40, 000 causal interaction effects. See **Online methods** for full details of phenotype construction.

**Figure S4:**
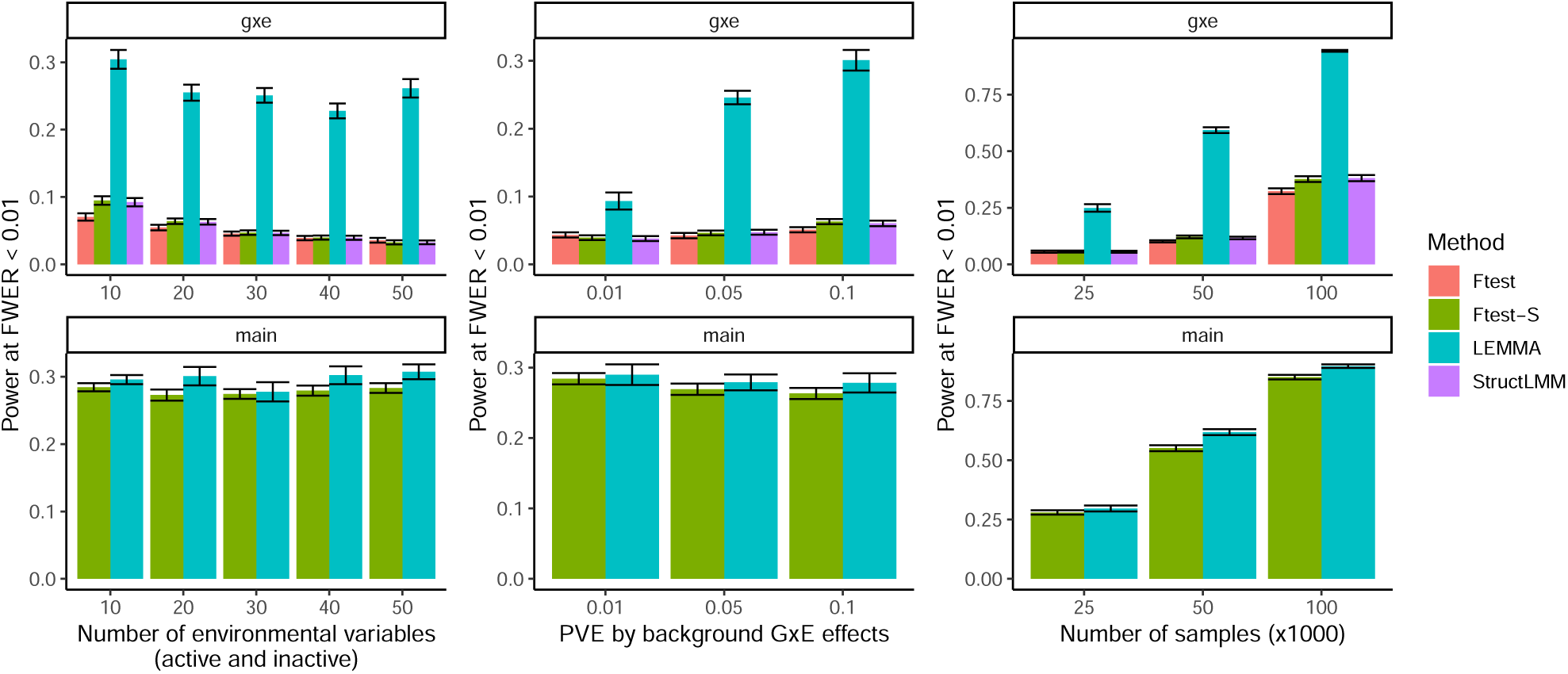
Power to detect causal SNPs in simulation. Power to detect SNP GxE interaction effects (top) and SNP main effects (bottom), whilst varying (a) the number of environmental variables, (b) proportion of trait variance explained by background GxE effects and (c) sample size. Power was assessed as the proportion of 60 causal SNPs detected at *p* < 0.01 (Family Wise Error Rate; FWER < 0.01), where causal SNPs main and GxE interaction effects each explained 0.00016% of trait variance. Simulations used genotypes sub-sampled from the UK Biobank and by default contained *N* = 25*K* samples, *M* = 100*K* SNPs, 6 environmental variables that contributed to the ES and 24 that did not (default parameters denoted by stars). We performed 20 repeats for each scenario. See **Online methods** for full details of phenotype construction.

**Figure S5:**
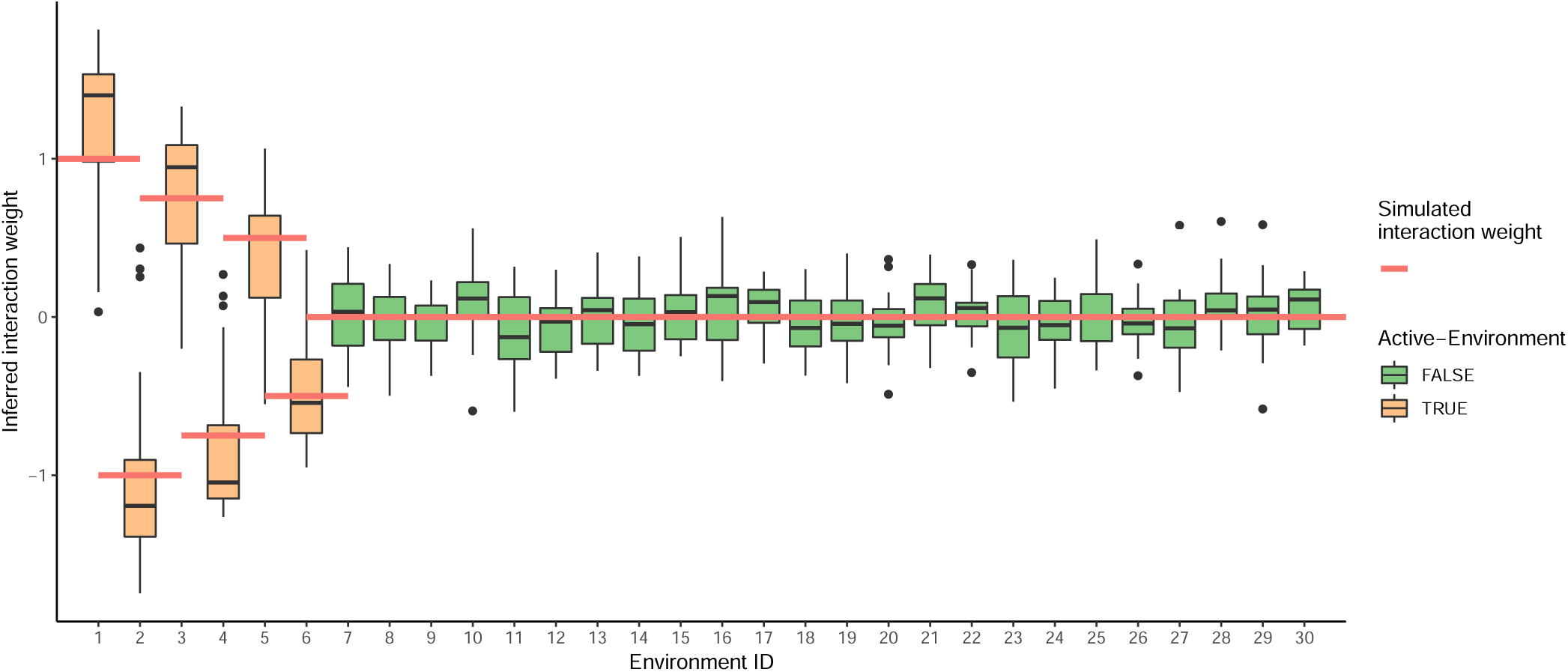
Estimation of ES weights in simulation. Boxplots of the environmental score (ES) weights estimated by LEMMA (left) over 20 simulations. Red lines denote true weights used to construct the simulated ES. Simulations performed with *N* = 25*k* samples, *M* = 100*k* SNPs and *L* = 30 environments (of which 6 were active). Phenotypes were constructed with *M*_causal, main-effects_ = 5000 SNPs explaining 20% of trait variance and *M*_causal, GxE-effects_ = 2500 SNPs explaining 5% of trait variance. LEMMA is invariant to a sign change in both the interaction weights and interaction SNP effects, so ES weights are automatically re-scaled such that the largest weight is positive before plotting.

**Figure S6:**
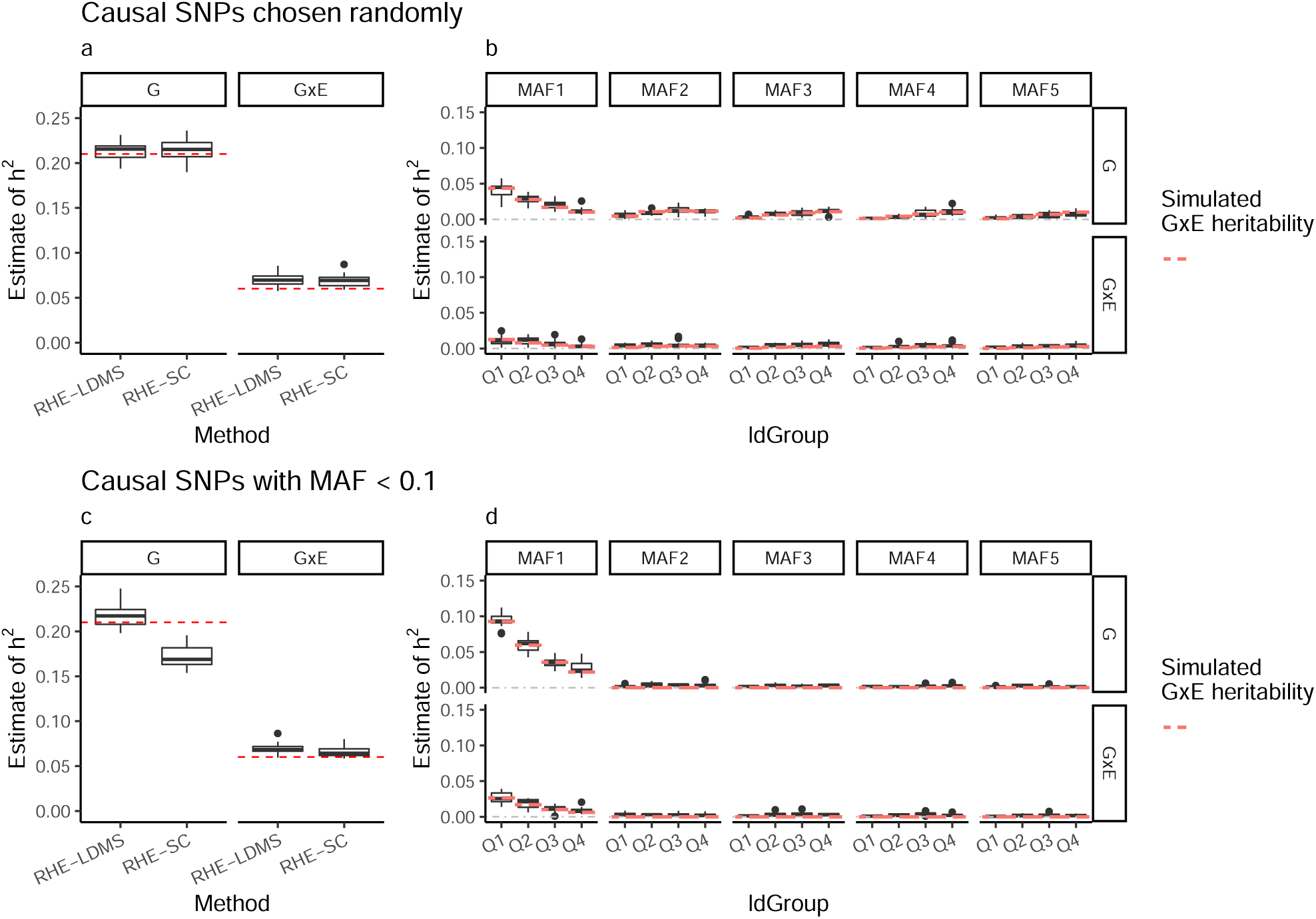
Heritability estimates stratified by LD and MAF in simulation. Comparison of heritability estimates using RHE-SC and RHE-LDMS when causal SNPs were drawn (a) at random or (c) only from low frequency (MAF < 0.1) SNPs. Heritability estimates (using RHE-LDMS) stratified by MAF when causal SNPs were drawn (b) at random or (d) only from low frequency (MAF < 0.1) SNPs. Simulations performed with *N* = 25*K* samples, *M* = 100*K* SNPs and the default simulation parameters described in **Online Methods**. Abbreviations; MAF, minor allele frequency; RHE-SC, randomized HE-regression with a single SNP component^23^; RHE-LDMS, multi-component randomized HE-regression^24^.

**Figure S7:**
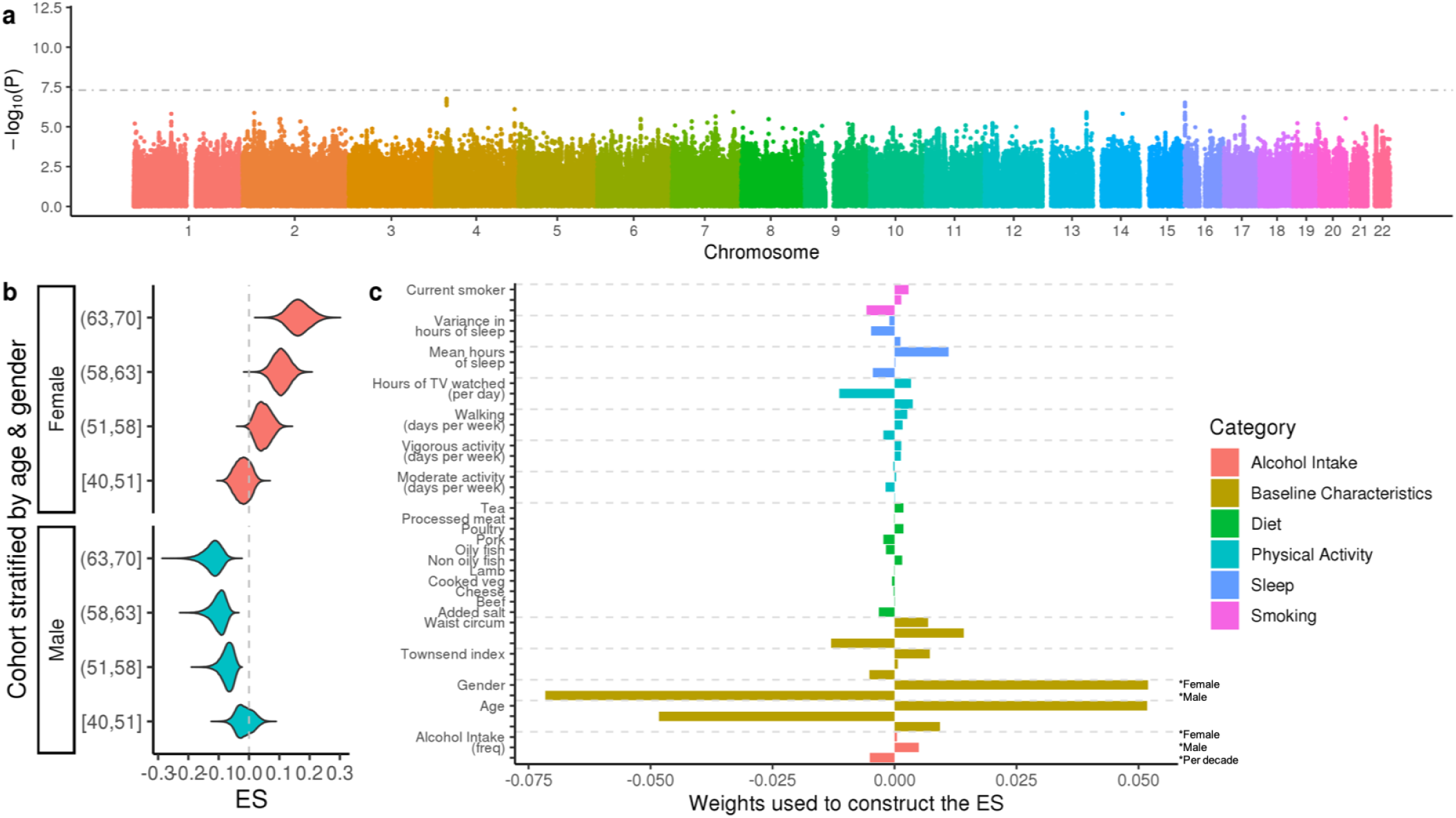
GxE analysis of PP in the UK Biobank. (a) LEMMA association statistics testing for multiplicative GxE interactions at each SNP. The horizontal grey line denotes (*p* = 5 × 10^−8^), *p*-values are shown on the − log_10_ scale. (b) Distribution of the environmental score (ES), stratified by gender and age quantile. (c) Weights used to construct the ES. Dietary variables have a single weight shown on the per standard deviation (s.d) scale. ‘Gender’ has two weights; a gender specific intercept for women (first) and men (second). Remaining non-dietary variables have three weights; (first) a per s.d effect for women only, (second) a per s.d effect for men only, (third) a per s.d per decade effect which is the same for both genders. s.d for the male and female specific weights is computed for each gender separately. Age is computed as the number of decades aged from 40. See **Online Methods** for details.

**Figure S8:**
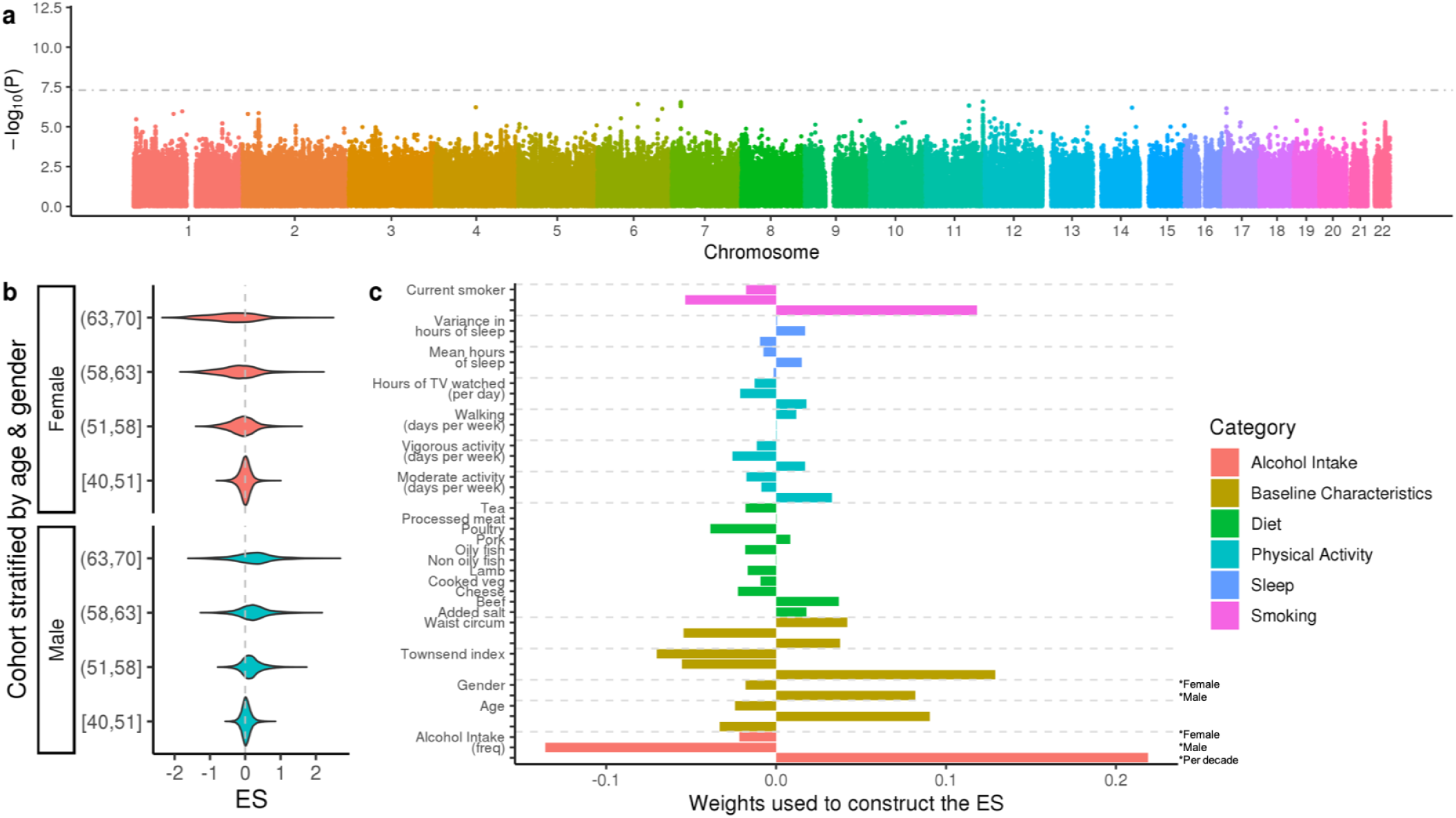
GxE analysis of SBP in the UK Biobank. (a) LEMMA association statistics testing for multiplicative GxE interactions at each SNP. The horizontal grey line denotes (*p* = 5 × 10^−8^), *p*-values are shown on the − log_10_ scale. (b) Distribution of the environmental score (ES), stratified by gender and age quantile. (c) Weights used to construct the ES. Dietary variables have a single weight shown on the per standard deviation (s.d) scale. ‘Gender’ has two weights; a gender specific intercept for women (first) and men (second). Remaining non-dietary variables have three weights; (first) a per s.d effect for women only, (second) a per s.d effect for men only, (third) a per s.d per decade effect which is the same for both genders. s.d for the male and female specific weights is computed for each gender separately. Age is computed as the number of decades aged from 40. See **Online Methods** for details.

**Figure S9:**
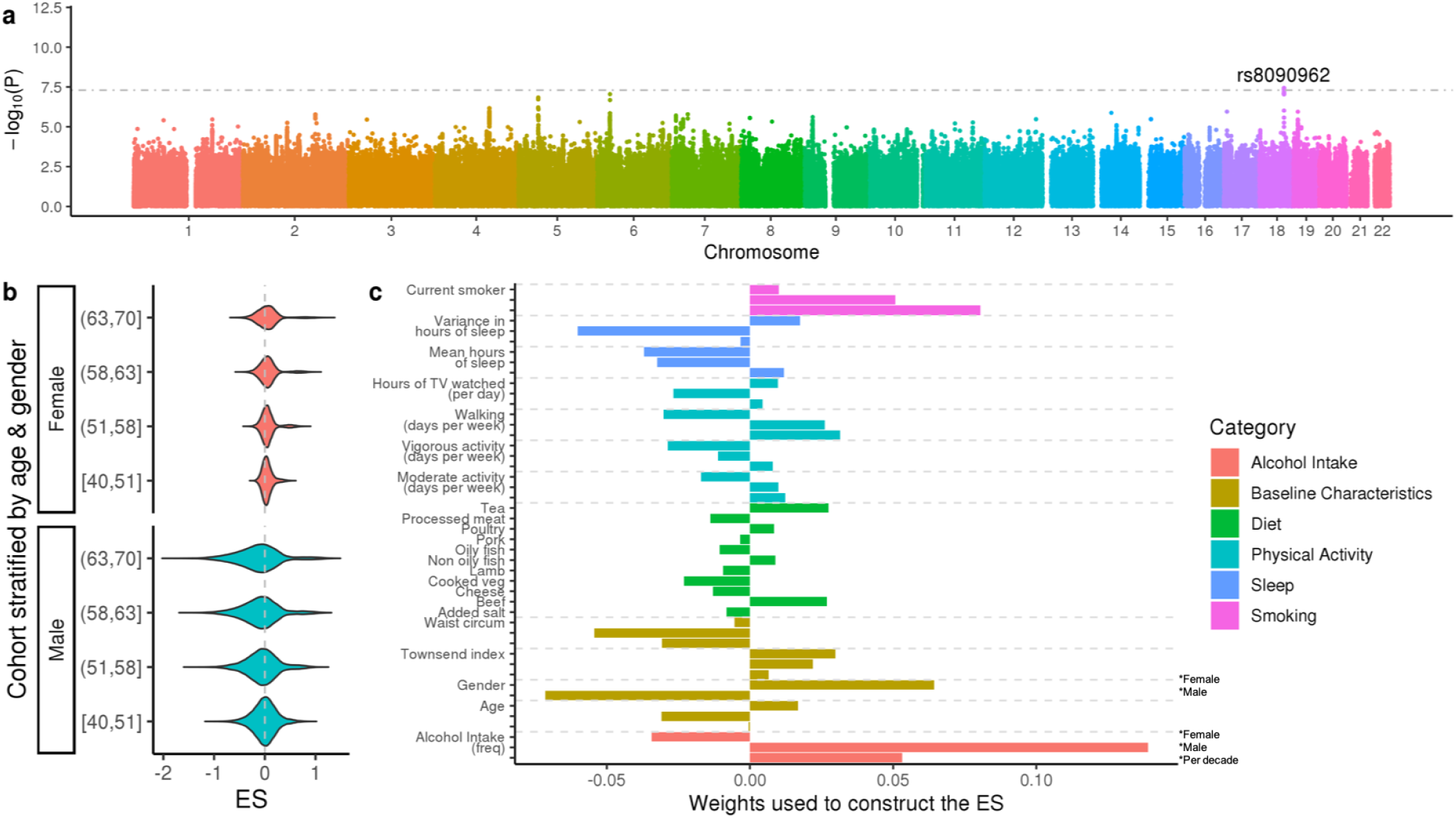
GxE analysis of DBP in the UK Biobank. (a) LEMMA association statistics testing for multiplicative GxE interactions at each SNP. The horizontal grey line denotes (*p* = 5 × 10^−8^), *p*-values are shown on the − log_10_ scale. (b) Distribution of the environmental score (ES), stratified by gender and age quantile. (c) Weights used to construct the ES. Dietary variables have a single weight shown on the per standard deviation (s.d) scale. ‘Gender’ has two weights; a gender specific intercept for women (first) and men (second). Remaining non-dietary variables have three weights; (first) a per s.d effect for women only, (second) a per s.d effect for men only, (third) a per s.d per decade effect which is the same for both genders. s.d for the male and female specific weights is computed for each gender separately. Age is computed as the number of decades aged from 40. See **Online Methods** for details.

**Figure S10:**
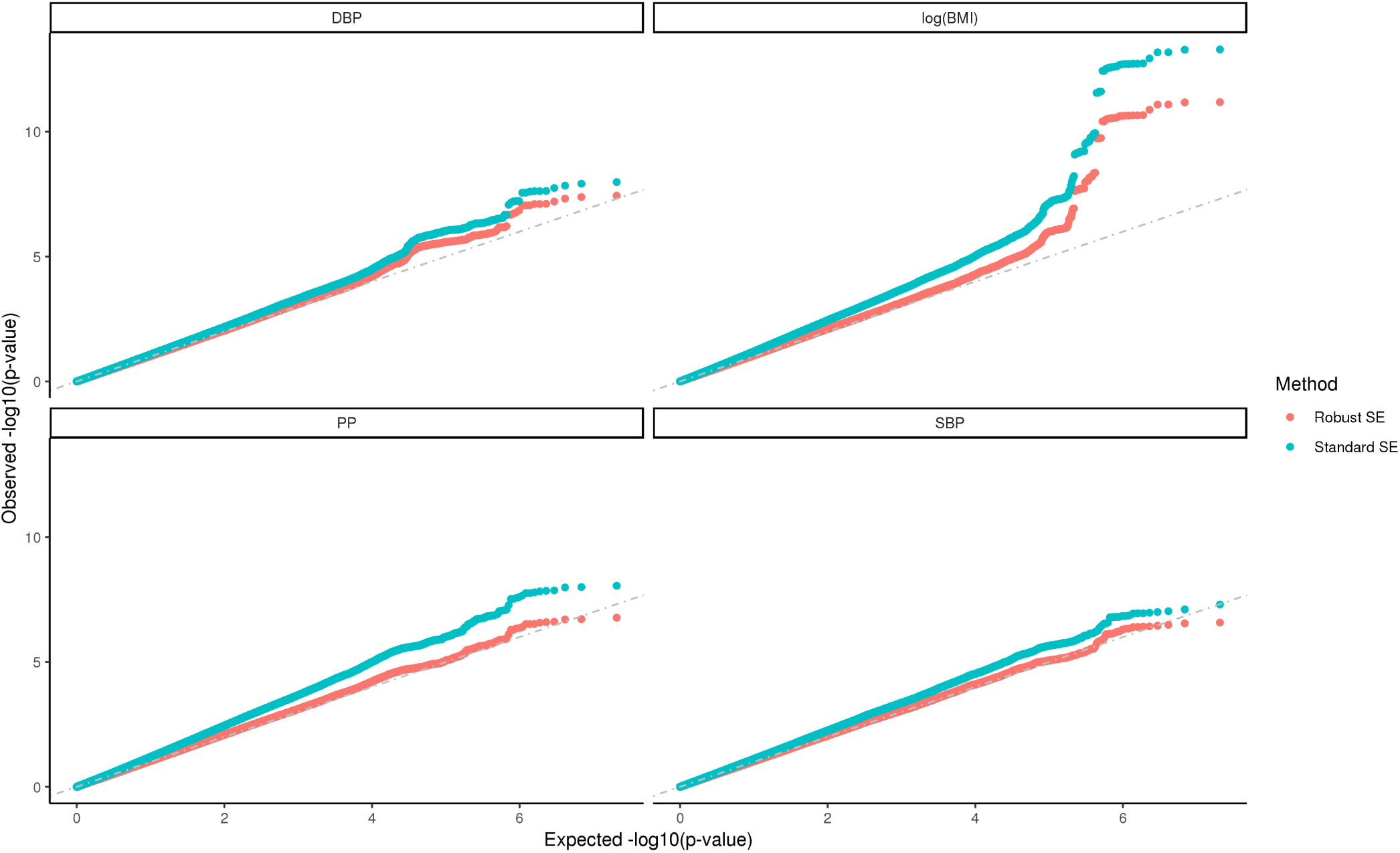
Effect of using robust standard errors for GxE interaction tests in the UK Biobank. QQ plots of the observed LEMMA − log_10_(*p*) values for GxE interactions at imputed SNPs for four UK Biobank traits, with and without robust standard errors. The grey dotted line denotes expected − log_10_(*p*)-values under a null model. Association tests using ‘Robust’ standard errors are well calibrated in both homoskedastic and heteroskedastic regimes (see **Online Methods**) and are used in all follow up analysis. Genomic control statistics were 1.275, 1.271, 1.163, 1.111 for logBMI, PP, SBP and DBP respectively using homoskedastic standard errors and 1.062, 1.047, 1.037, 1.027 for logBMI, PP, SBP and DBP respectively using robust standard errors.

**Figure S11:**
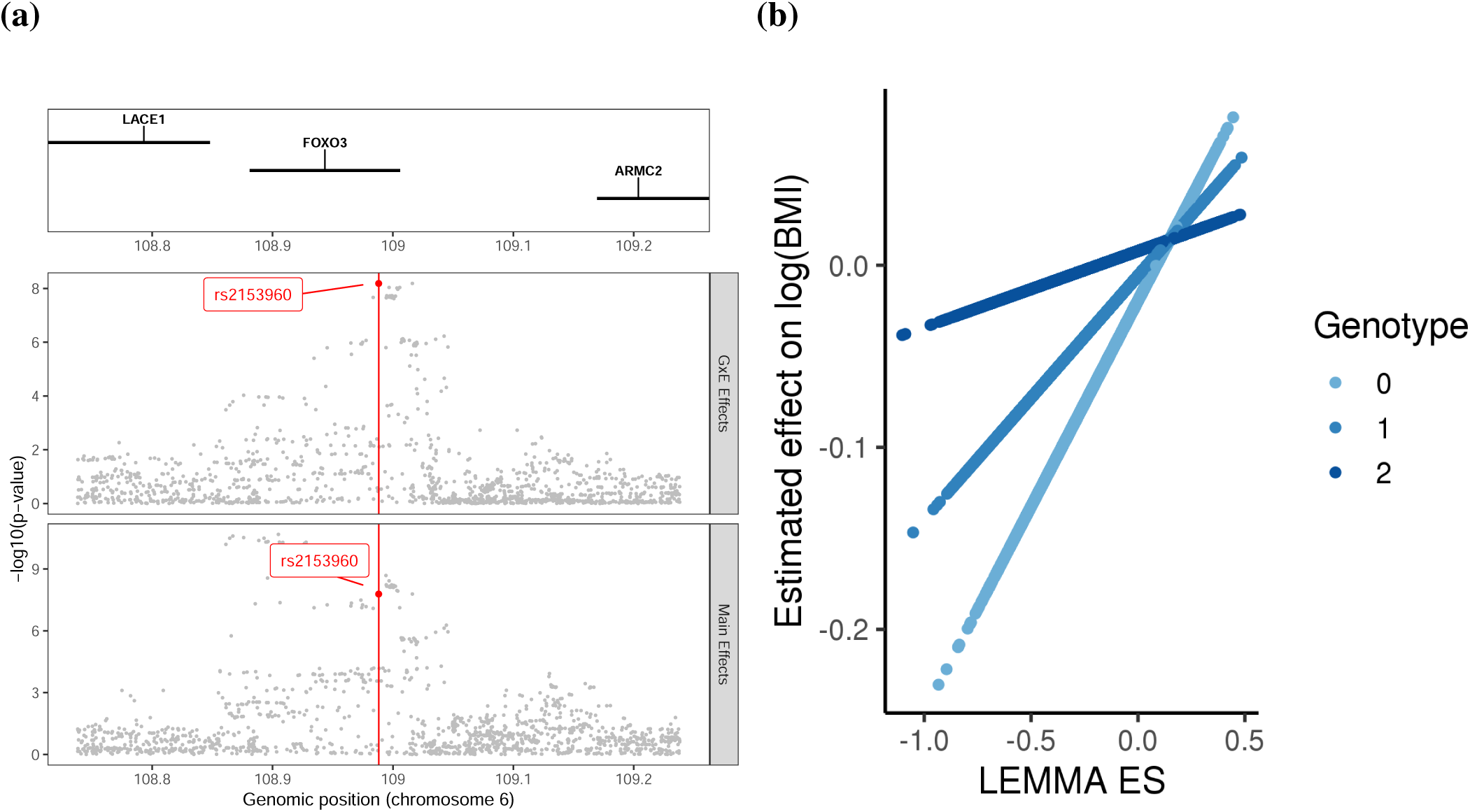
Estimated GxE effect rs2153960 on logBMI. (a) Regional plot of the main and interaction effects of SNPs within 250KB of rs2153960, (b) the estimated effect of rs2153960 on logBMI as a function of the environmental score (ES).

**Figure S12:**
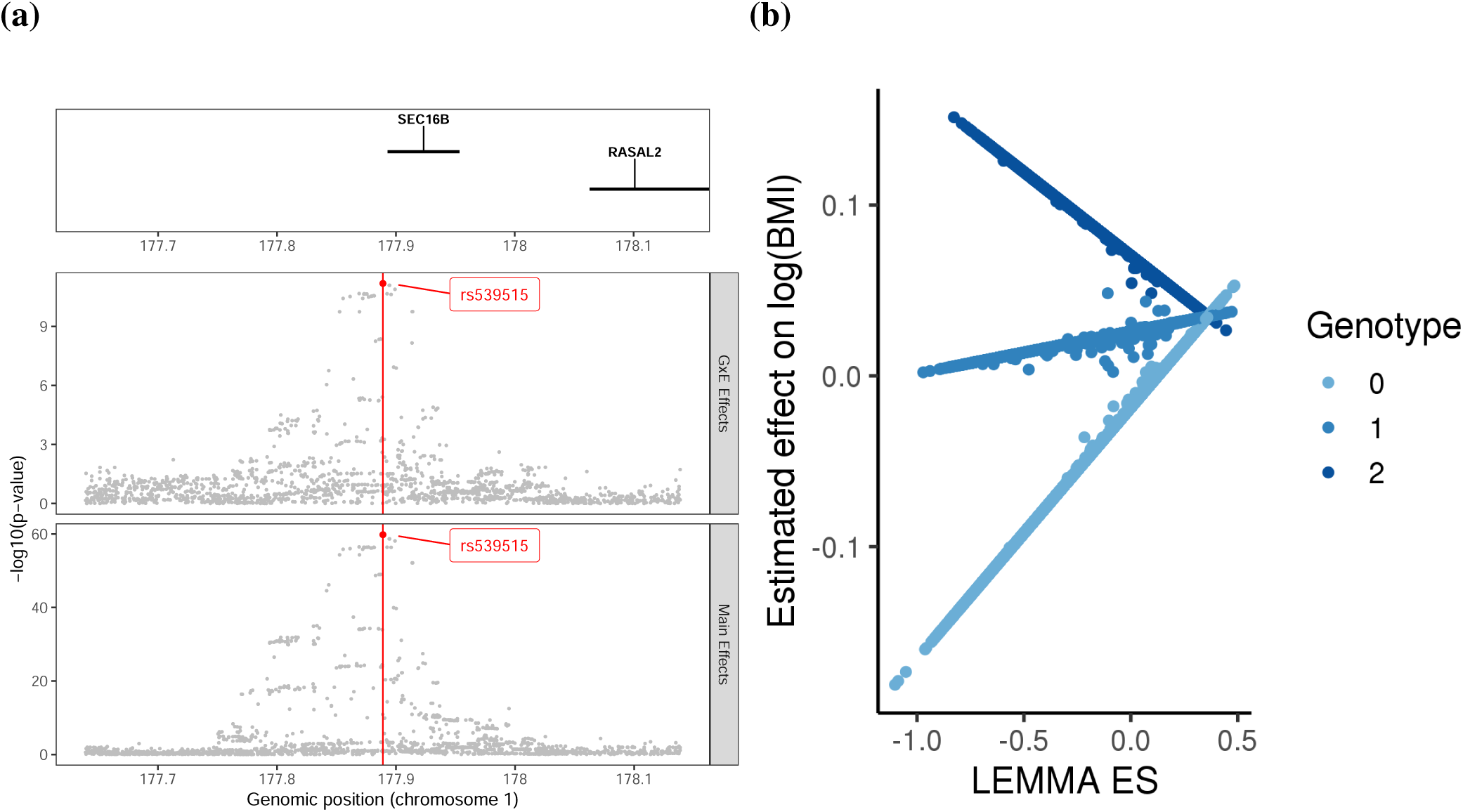
Estimated GxE effect rs539515 on logBMI. (a) Regional plot of the main and interaction effects of SNPs within 250KB of rs539515, (b) the estimated effect of rs539515 on logBMI as a function of the environmental score (ES).

**Figure S13:**
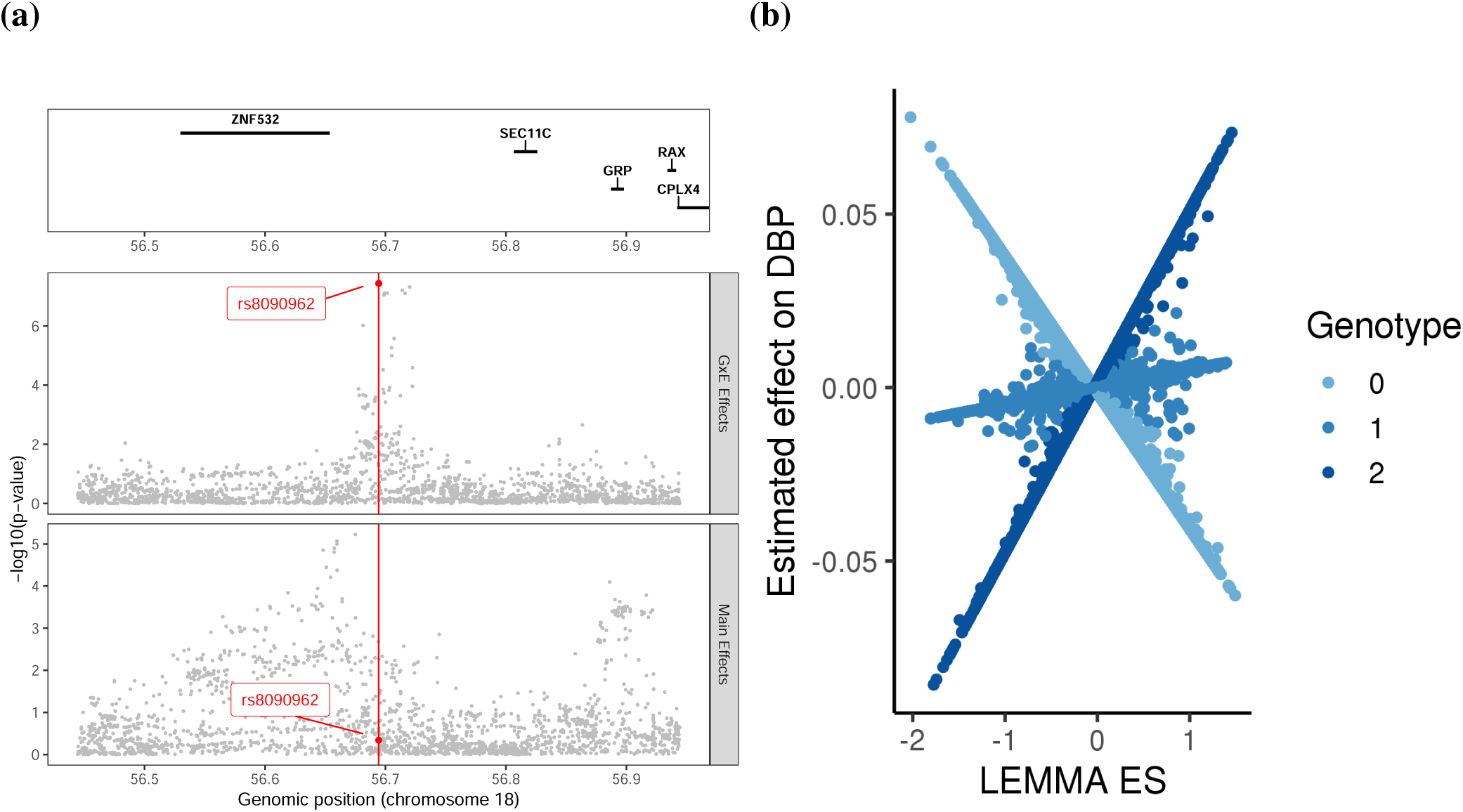
Estimated GxE effect rs8090962 on DBP. (a) Regional plot of the main and interaction effects of SNPs within 250KB of rs8090962, (b) the estimated effect of rs8090962 on DBP as a function of the environmental score (ES).

**Figure S14:**
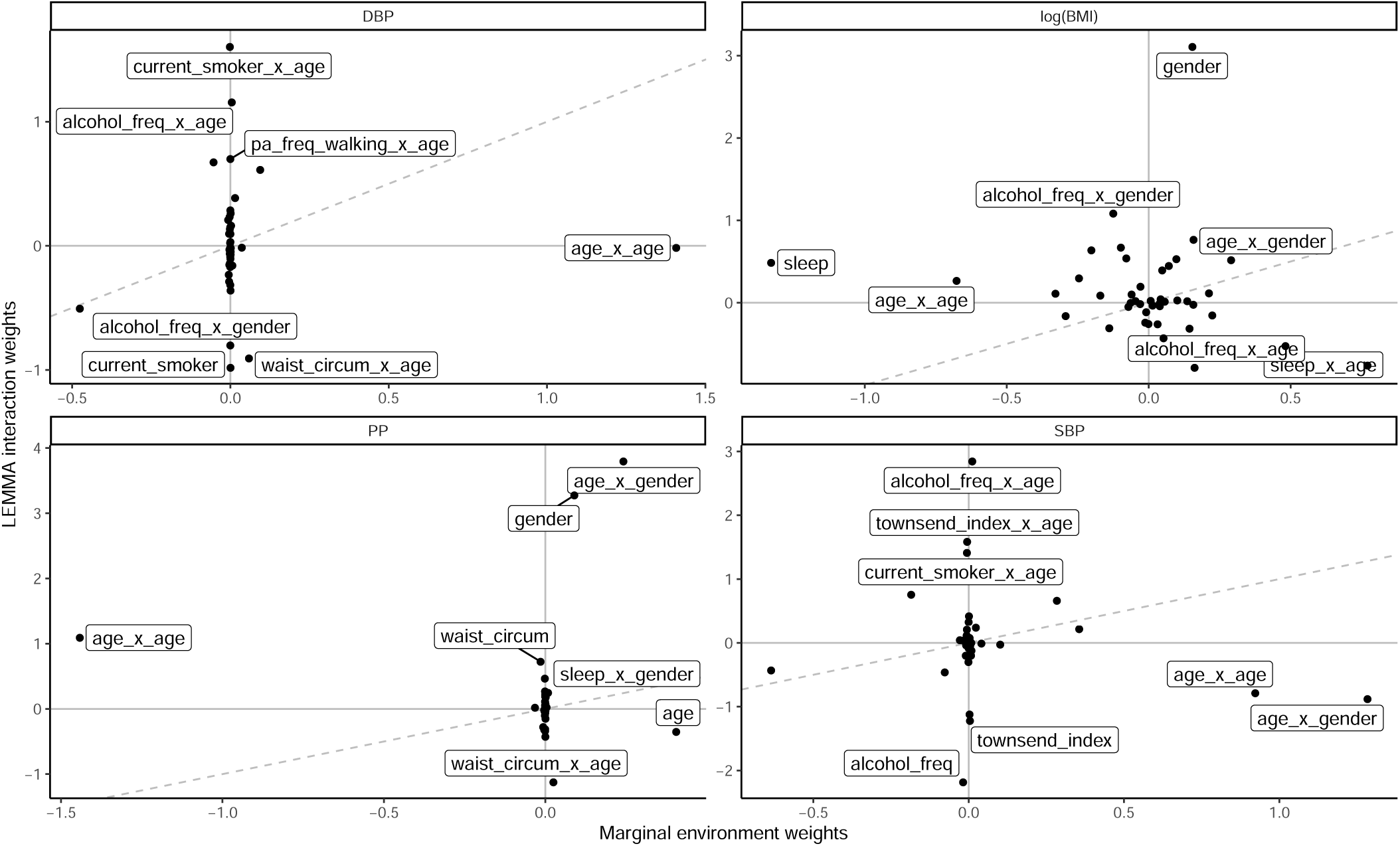
Comparison of the LEMMA vs marginal environmental score. Interaction weights of the marginal environmental score were estimated from multivariate linear regression, using all the non-genetic covariates used by LEMMA. Interactions weights were all rescaled so that the corresponding ES had variance one. The dashed grey line represents the *y* = *x* line.

**Figure S15:**
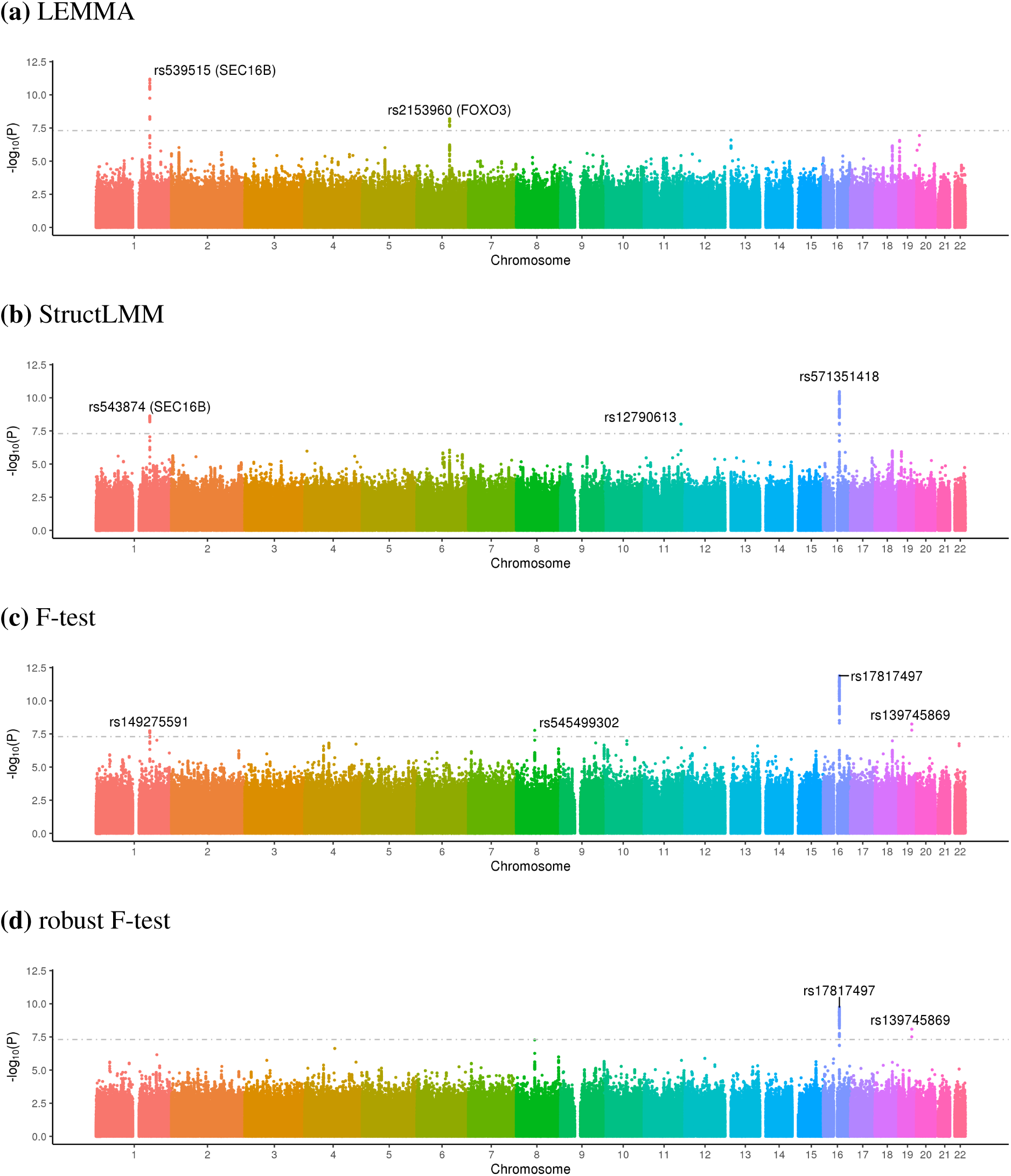
GxE association statistics for logBMI. Manhattan plots displaying the negative log_1_0 *p* values from GxE interaction tests at 10, 295, 038 imputed SNPs applied to logBMI in the UK Biobank. GxE interaction tests were computed using (a) LEMMA, (b) StructLMM, (c) the F-test and (d) the robust F-test. The horizontal grey line denotes (*p* = 5 × 10^−8^).

**Figure S16:**
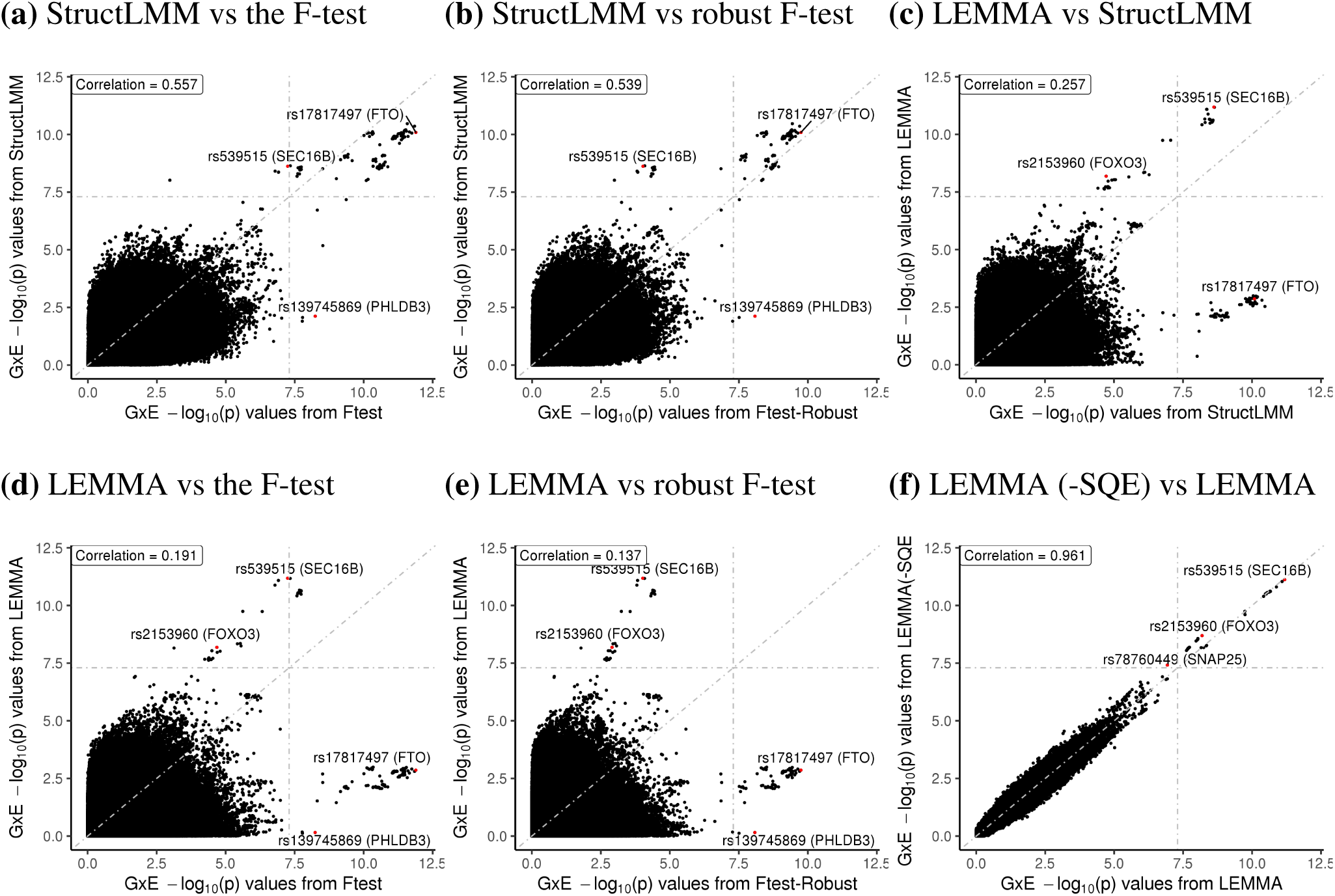
Comparison of GxE association statistics for logBMI. Comparison of negative log_10_ *p* values obtained from LEMMA, StructLMM, the F-test and the robust F-test in an analysis of logBMI in the UK Biobank. Grey lines denote (*p* = 5 × 10^−8^) and the *y* = *x* axis. Pearson correlation is shown in a label at the top left of each plot. Red points denote the sentinel SNP for each locus.

**Figure S17:**
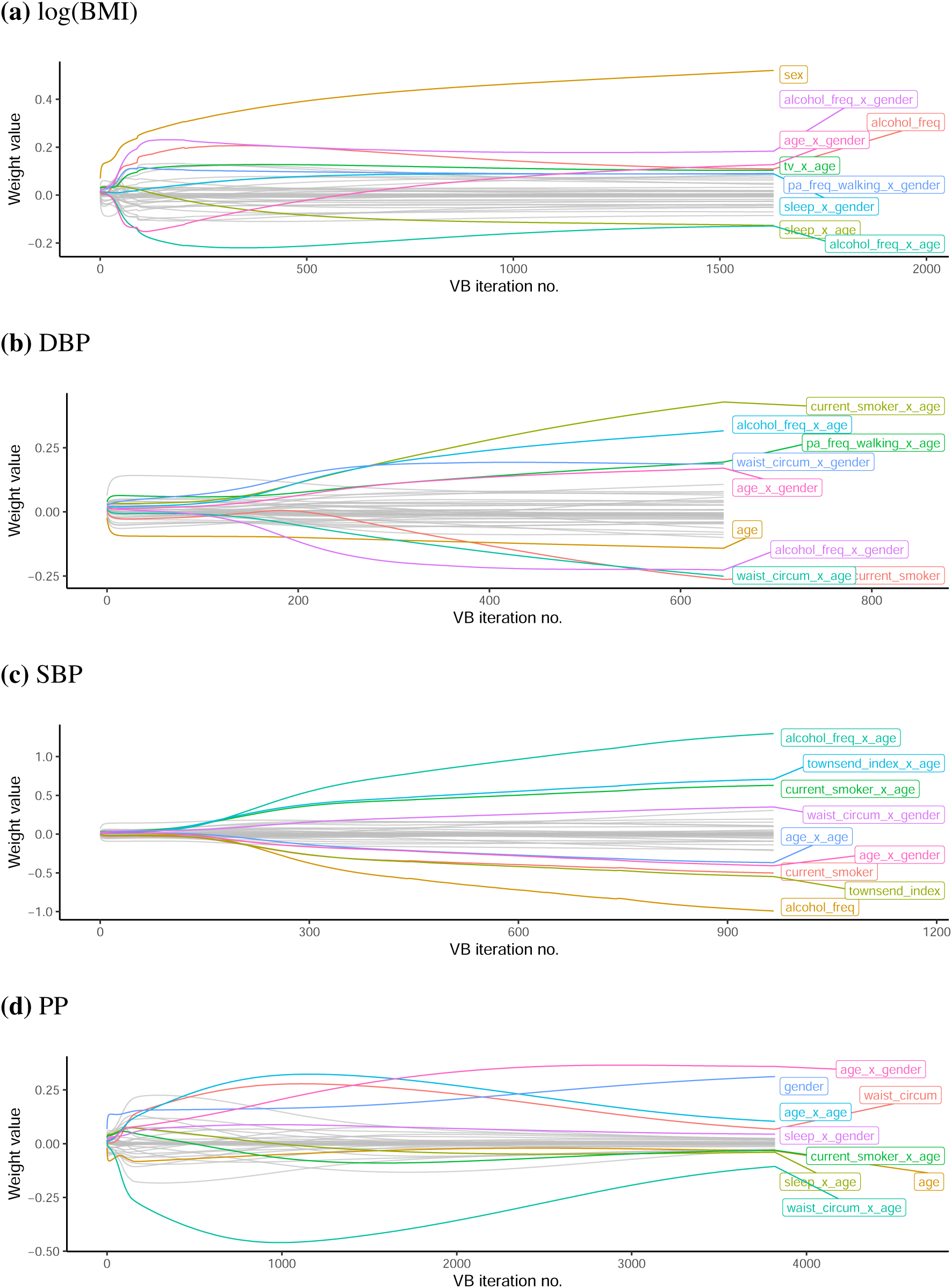
Inference of environmental score weights from GxE analyses of four quantitative traits in the UK Biobank. Evolution of the environmental score weights as LEMMA performs successive passes through the data.

**Figure S18:**
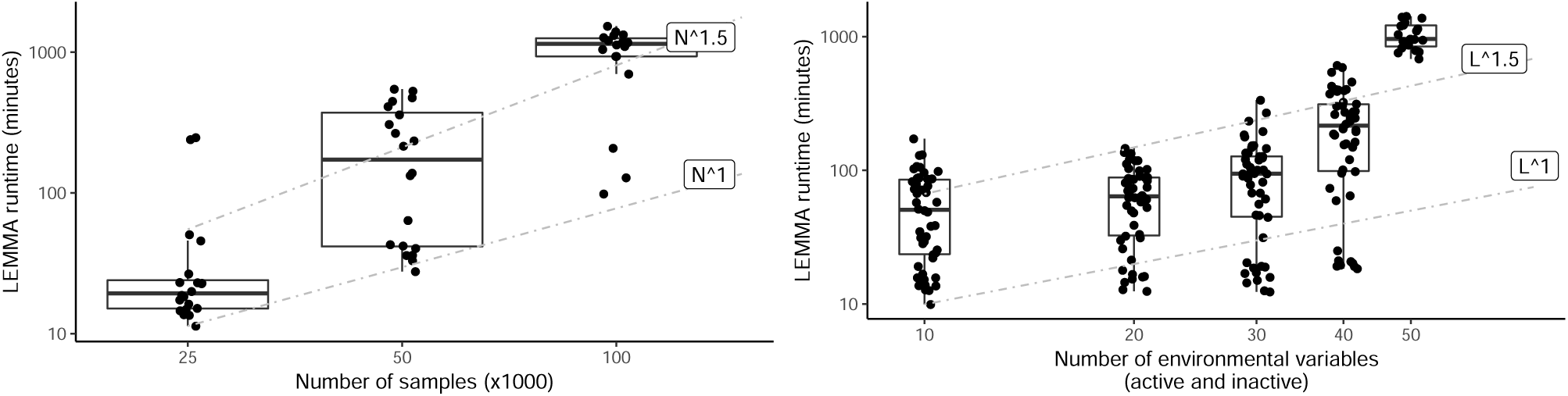
Log-log plots showing runtime of the variational bayes algorithm used to perform whole genome regression by LEMMA, as a function of sample size (left) and the number of environmental variables (right). Unless otherwise stated simulations were performed using *N* = 25*k* samples, *M* = 100*k* SNPs and *L* = 30 environmental variables. Phenotypes were constructed using 2500 non-zero main effects explaining 20% of variance, 1250 nonzero interaction effects explaining 5% of variance and 6 active environmental variables. See **Online methods** for full details of phenotype construction.

**Figure S19:**
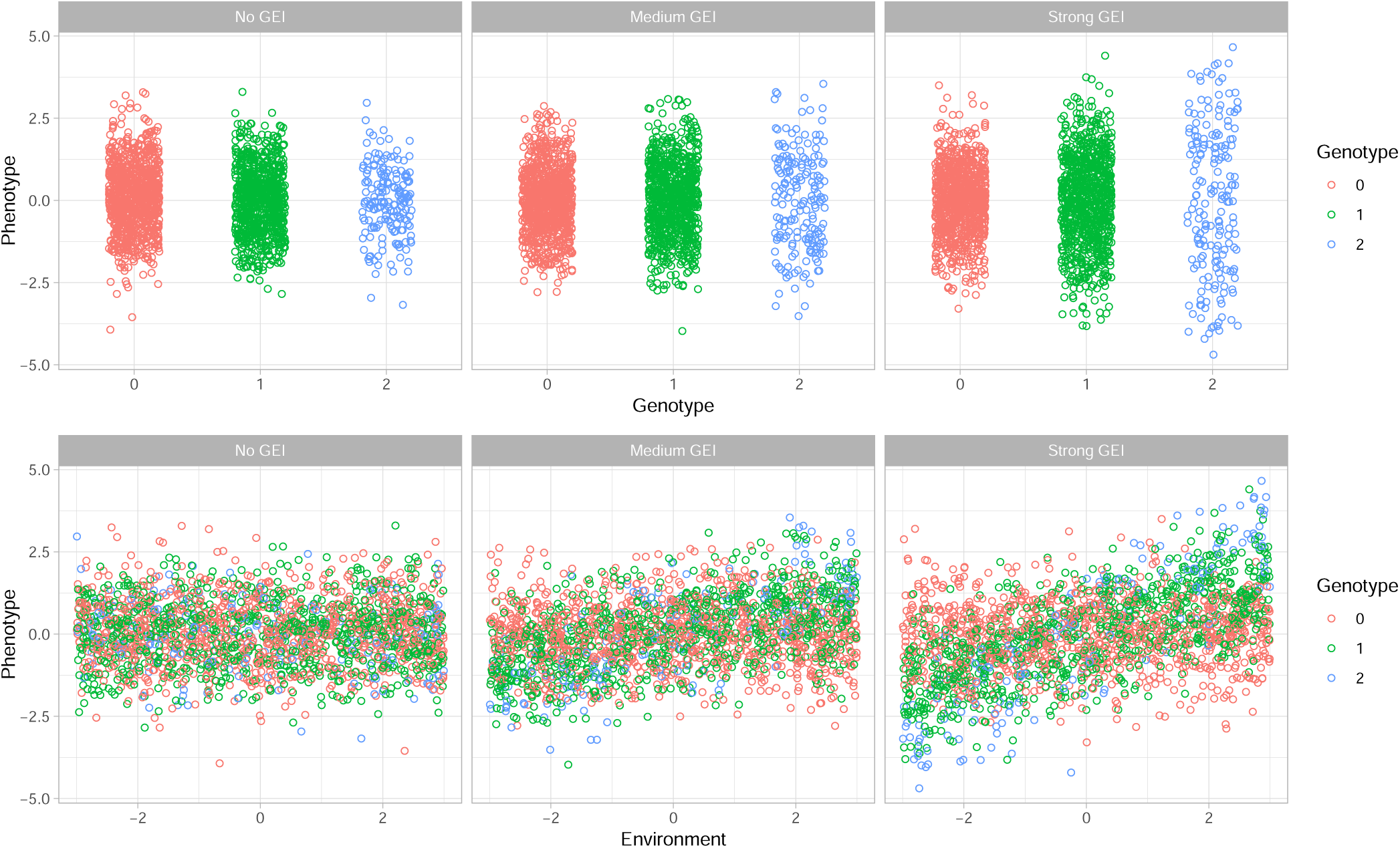
Visualization of differences in variance induced by a multiplicative Gene-x-Environment effect. Differences in phenotypic variance by genotype group (top) and by strength of the environmental exposure (bottom). The phenotype was simulated using 2000 individuals on the basis of a multiplication interaction between a single genotype (minor allele frequency 0.3) and an environment (uniformly distributed over [−2, 2]). From left to right; the GxE interaction explained 0%, 20%, 40% of trait variance.

**Table S1:** Quality control and time to convergence of the WGR analyses. Time for the whole genome regression analysis to converge is reported for four quantitative traits in the UK Biobank, as well as the number of SNPs and samples passing quality control and the number of covariates controlled for. ‘Other covariates’ consisted of the top 20 genetic principal components as reported by the UK Biobank, age^3^, age^2^× gender, age^3^× gender, a binary indicator for the genotype chip and (for blood pressure traits only) BMI. Environmental variables used (including lower orders of age and gender) are described in (**Online Methods**). To control for potential bias due to non-linear dependence between the phenotype and heritable environmental variables, we tested each environmental variable and included any significant squared effects as additional covariates (**Online Methods**) *based on the average per-iteration cost of 243 seconds, using 32 cores distributed across a cluster with Xeon E5-2667 v4 3.2Ghz processors.

| Trait | No. samples | No. SNPs | No. envs | No. envs-sq | No. other covars | No. covars total | Iterations for WGR converge | Time for WGR to converge* |
| --- | --- | --- | --- | --- | --- | --- | --- | --- |
| log(BMI) | 281149 | 642095 | 42 | 30 | 24 | 96 | 1631 | 78 hours 18 mins |
| PP | 280749 | 642102 | 45 | 13 | 25 | 83 | 3821 | 183 hours 26 mins |
| SBP | 280749 | 642102 | 45 | 15 | 25 | 85 | 967 | 46 hours 25 mins |
| DBP | 280749 | 642102 | 45 | 15 | 25 | 85 | 646 | 31 hours 00 mins |

**Table S2:** Partitioned heritability estimates for four quantitative traits in the UK Biobank. Comparison of the heritability estimates obtained using genotyped SNPs with RHE-SC, genotyped SNPs with RHE-LDMS, and common imputed SNPs (MAF> 0.01 in the full UK Biobank cohort) with RHE-LDMS. GxE heritability estimates were were obtained using the ES from each model fit. All analyses controlled for the same covariates used in the WGR analysis (including the top 20 principal components). Abbreviations; s.e, standard error estimated using the block jack-knife (see **Online Methods**); 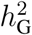, heritability due to additive genetic effects; 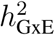, heritability due to multiplicative GxE effects; RHE, randomized HE-regression^23, 24^; SC, single SNP component; LDMS, SNPs stratified by minor allele frequency and LDscore (20 components).

| Trait | Genotyped (SC) |  | Genotyped (LDMS) |  | Common Imputed (LDMS) |  |
| --- | --- | --- | --- | --- | --- | --- |
| | $h_G^2$ (s.e) | $h_{GxE}^2$ (s.e) | $h_G^2$ (s.e) | $h_{GxE}^2$ (s.e) | $h_G^2$ (s.e) | $h_{GxE}^2$ (s.e) |
| log BMI | 0.259 (0.069) | 0.071 (0.009) | 0.237 (0.126) | 0.086 (0.024) | 0.274 (0.056) | 0.093 (0.028) |
| PP | 0.233 (0.039) | 0.075 (0.018) | 0.203 (0.084) | 0.111 (0.021) | 0.228 (0.051) | 0.125 (0.028) |
| SBP | 0.24 (0.053) | 0.033 (0.003) | 0.223 (0.095) | 0.038 (0.017) | 0.251 (0.05) | 0.039 (0.023) |
| DBP | 0.277 (0.034) | 0.014 (0.001) | 0.231 (0.079) | 0.016 (0.017) | 0.254 (0.05) | 0.016 (0.02) |

**Table S5:**
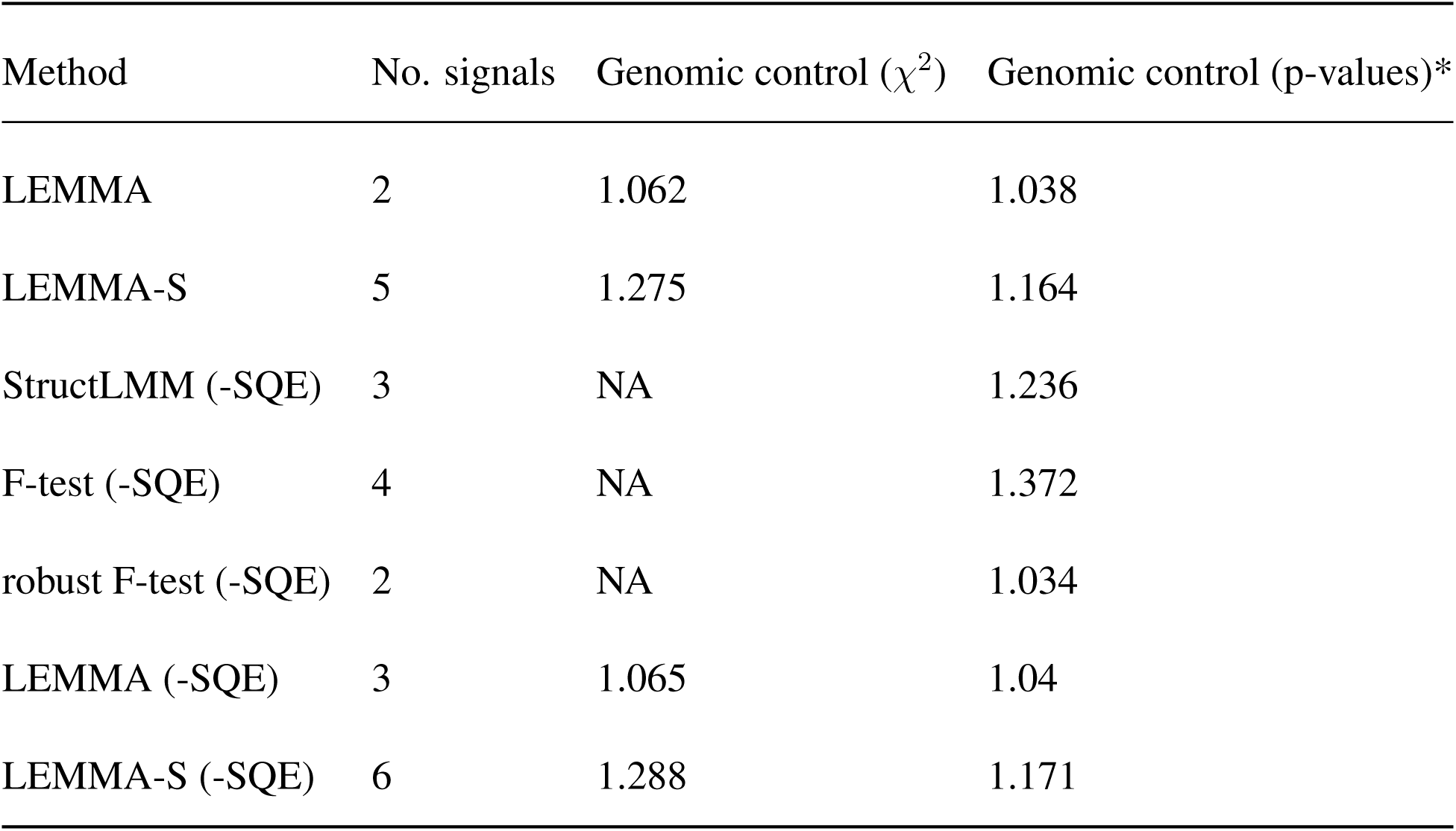
Comparison of the number of genome-wide significant GxE associations and genomics control statistics from a GxE analysis of logBMI in the UK Biobank. The number of independent loci (at least 0.5cM apart) with genome-wide significant GxE interaction effects and genomic control statistics for seven different methods applied to logBMI in the UK Biobank. Genomic control is computed from GxE interaction tests statistics from 10, 295, 038 imputed SNPs. Abbreviations; LEMMA-S, LEMMA with a homoskedastic test statistic (see **Online Methods**); (-SQE), significant squared environmental variables (Bonferroni correction) not included as additional covariates. *The test statistics from StructLMM, F-test and the robust F-test are not 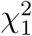 distributed. Hence for these methods we use *λ*_GC_ = log_10_(*m*)*/* log_1_ 0(0.5), where *m* is the median *p*-value, to denote the genomic control statistic as suggested by Moore *et al.*^7^.

| Method | No. signals | Genomic control ($\chi^2$) | Genomic control (p-values)* |
| --- | --- | --- | --- |
| LEMMA | 2 | 1.062 | 1.038 |
| LEMMA-S | 5 | 1.275 | 1.164 |
| StructLMM (-SQE) | 3 | NA | 1.236 |
| F-test (-SQE) | 4 | NA | 1.372 |
| robust F-test (-SQE) | 2 | NA | 1.034 |
| LEMMA (-SQE) | 3 | 1.065 | 1.04 |
| LEMMA-S (-SQE) | 6 | 1.288 | 1.171 |

**Table S7:** Association between genetic principle components and the environmental score for four traits in the UK Biobank. Associations computed using ordinary least squares to regress the environmental score against the top 20 principle components (with an intercept included). Association strength reported using negative log_10_(*P*)-values from a standard t-test. Abbreviations; PC, genetic principle component; ES, environmental score.

|  | logBMI ES | PP ES | SBP ES | DBP ES |
| --- | --- | --- | --- | --- |
| PC1 | 2.182 | 0.023 | 0.467 | 0.403 |
| PC2 | 0.254 | 0.153 | 0.197 | 0.088 |
| PC3 | 0.022 | 0.382 | 0.405 | 0.295 |
| PC4 | 0.657 | 0.465 | 0.091 | 0.095 |
| PC5 | 15.075 | 2.652 | 68.293 | 71.439 |
| PC6 | 0.703 | 0.780 | 0.020 | 0.480 |
| PC7 | 0.469 | 0.095 | 0.933 | 0.998 |
| PC8 | 0.778 | 1.898 | 2.784 | 1.848 |
| PC9 | 0.675 | 0.878 | 14.130 | 27.305 |
| PC10 | 0.814 | 0.081 | 0.647 | 0.276 |
| PC11 | 2.759 | 0.255 | 6.547 | 7.970 |
| PC12 | 0.319 | 0.974 | 0.486 | 0.511 |
| PC13 | 0.659 | 0.508 | 0.475 | 2.003 |
| PC14 | 3.301 | 5.779 | 2.575 | 4.454 |
| PC15 | 0.186 | 0.418 | 0.300 | 0.531 |
| PC16 | 3.554 | 3.141 | 2.830 | 6.560 |
| PC17 | 0.695 | 1.590 | 1.208 | 0.061 |
| PC18 | 3.965 | 0.215 | 0.681 | 0.884 |
| PC19 | 0.350 | 0.375 | 0.099 | 0.409 |
| PC20 | 2.463 | 1.208 | 1.228 | 0.720 |

**Table S8:** Sensitivity of partitioned heritability estimates to ES-x-PCs interaction in the UK Biobank. Heritability estimates were computed using genotyped SNPs with RHE-LDMS and the ES from each WGR analysis. Left; heritability estimates obtained whilst controlling for the same covariates used in the WGR analysis (including the top 20 principal components), right; heritiability estimates obtained whilst additionally controlling for multiplicative interactios between the ES and genetic PCs. Abbreviations; s.e, standard error estimated using the block jackknife (see **Online Methods**); 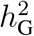, heritability due to additive genetic effects; 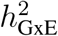, heritability due to multiplicative GxE effects; RHE, randomized HE-regression^23, 24^; LDMS, SNPs stratified by minor allele frequency and LDscore (20 components).

| Trait | Genotyped (LDMS) |  | Genotyped (LDMS), additionally controlling for ES-x-PCs |  |
| --- | --- | --- | --- | --- |
| | $h_G^2$ (s.e) | $h_{G \times E}^2$ (s.e) | $h_G^2$ (s.e) | $h_{G \times E}^2$ (s.e) |
| log BMI | 0.2366 (0.1259) | 0.0862 (0.0237) | 0.2368 (0.1259) | 0.0862 (0.0236) |
| PP | 0.2025 (0.0841) | 0.1110 (0.0208) | 0.2025 (0.0840) | 0.1113 (0.0208) |
| SBP | 0.2225 (0.0954) | 0.0377 (0.0168) | 0.2226 (0.0955) | 0.0360 (0.0166) |
| DBP | 0.2308 (0.0793) | 0.0157 (0.0166) | 0.2308 (0.0793) | 0.0142 (0.0163) |

**Table S9:** Correlation between main SNP effects and interaction SNP effects. Correlation between main SNP effects and interaction SNP effects learnt during the LEMMA variational algorithm. Absolute correlation is used as *γ* is invariant to being multiplied by −1 (as LEMMA would apply the same transform to the ES).

| Trait | $\text{cor}(\text{ES}, \text{ES}_{\text{main}})$ | $\text{abs}(\text{cor}(X\beta, X\gamma))$ | $\text{abs}(\text{cor}(\beta, \gamma))$ |
| --- | --- | --- | --- |
| log(BMI) | -0.062 | 0.13680 | 0.05810 |
| PP | -0.019 | 0.05306 | 0.03324 |
| SBP | -0.297 | 0.01741 | 0.00816 |
| DBP | -0.088 | 0.02464 | 0.00732 |

## Notes

### Competing Interest Statement

The authors have declared no competing interest.

